# Network modeling identifies vitamin B2 control of AIFM2 deficit-driven hepatocyte death in MASLD

**DOI:** 10.1101/2024.10.10.617610

**Authors:** Julian Weihs, Fatima Baldo, Alessandra Cardinali, Gehad Youssef, Katarzyna A. Ludwik, Nils Haep, Peter Tang, Pavitra Kumar, Cornelius Engelmann, Susanna Quach, Mijuna Meindl, Martin Kucklick, Susanne Engelmann, Bruno Chilian, Michael Rothe, Isabella Lurje, Linda Hammerich, Merrit Romeike, Prakash Ramachandran, Timothy J. Kendall, Jonathan A. Fallowfield, Ludovic Vallier, Harald Stachelscheid, Igor M. Sauer, Christian Hudert, Takanori Takebe, Frank Tacke, Philip Bufler, Namshik Han, Milad Rezvani

## Abstract

Drug discovery for metabolic dysfunction-associated steatotic liver disease (MASLD) remains difficult due to network effects. We developed *MetaLiver*, a public resource to identify druggable nodes within complex interactomes. It incorporates a MASLD-specific knowledge graph that improves bioactive compound contextualization while reducing computing time 18.8-fold. Layering progression-resolved bulk and hepatocyte-specific single-cell MASLD data onto this graph identified the vitamin B2-dependent flavoproteome (FlavProt) as an injury-sensitive network. Induced pluripotent stem cell-derived and primary human hepatocytes validated that flavin-cofactor replenishment mitigates FlavProt-dependent dysregulation, augmenting metabolic functions under cellular stress. Saturating flavin-cofactor availability blunted human immune cell activation. Clinically, a FlavProt expression score significantly correlated with poor outcomes in retrospectively stratified MASLD patients. We identified the oxidoreductase AIFM2 from this score and established experimentally that its injury-associated deficit causes hepatocyte death yet is controllable by flavin-cofactor availability. This study reveals the FlavProt as critical yet modifiable in MASLD and establishes *MetaLiver* for network discovery.

---

MASLD is the most common liver disease and is not governed by a single pathway^1^. Instead, multisystem metabolic dysregulation and inflammation mediate the progressive failure of molecular networks in the liver at the tissue, cellular, and subcellular levels^2^. This complexity hampers hypothesis generation and slows the translation of models into clinical interventions, exemplified by the limited number of approved therapies and their modest efficacy: Resmetirom resolves steatohepatitis in ∼20% and Semaglutide in ∼28% of MASLD patients^3,4^ compared to placebo. Thus, capturing and modelling of the disease network architecture may facilitate the discovery of new effective therapeutic strategies. Progress in machine learning (ML-) based drug prediction has created new opportunities for rational, data-driven network identification^5^. Multi-level *data oceans* such as biomedical knowledge graphs (KGs) are a powerful resource to contextualize drugs within biological networks, but are affected by publication biases^6^ and high computing power requirements. Contrarily, tailored analysis strategies for clinical data sets can focus on specific disease aspects, which, however, often operate in isolation and single modalities, limiting holistic MASLD drug prioritization^7^. A complementary strategy using multiple ML–driven frameworks, including biomedical KGs and disease-stage-stratified human omics, could identify predominant networks for therapeutic interception. Applying such a strategy may further enable the prioritization of high-likelihood candidates for experimental validation, overcoming conventional large library screenings. As drug discovery for multifactorial diseases remains constrained by a trade-off between physiological relevance and experimental scalability^8^, such prioritization may enable pre-clinical validation in models involving induced pluripotent stem cell (iPSC-) and primary cell-derived systems that offer higher fidelity than transformed cell lines^9^.

Here, we develop *MetaLiver*, a digital framework for network identification and drug prioritization in MASLD. *MetaLiver* identified the flavoproteome (FlavProt) network as critical and tractable to mitigate MASLD (**Fig. 1**). We first generated a disease-specific KG by expanding MASLD-relevant nodes and augmented this network using liver transcriptomics from a patient data set. We then identified progression-correlated metabolic and hepatocyte-specific network nodes from bulk and single-cell patient data, nominating the FlavProt as a key target network. As this network is poorly described in MASLD, we validated its dysregulation and therapeutic potential in iPSC-derived and primary human hepatocytes undergoing multiple compounded injuries (Ci-Heps). Replenishing the availability of vitamin B2-derived FlavProt cofactor flavin adenine dinucleotide (FAD) alone mitigated injury-associated network perturbations, on molecular and functional levels, under compounded metabolic stress. Finally, we identified a polygenic FlavProt expression score that accurately discriminated liver decompensation risk in MASLD patients. This FlavProt signature identified decreased AIFM2 activity as a contributor to MASLD-associated hepatocyte death, which we experimentally mitigated by FAD replenishment.

**Fig. 1:**
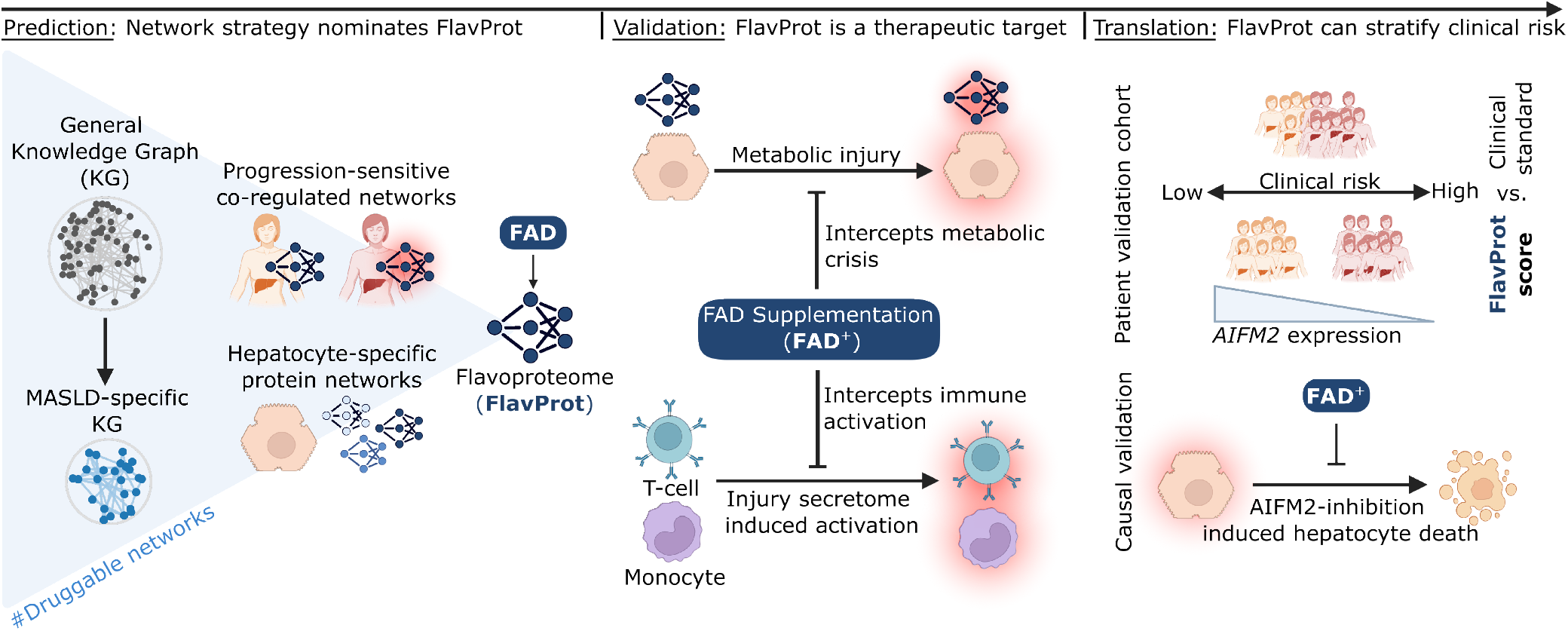
Strategy of integrating computational network models with functional experimentation ex vivo and retrospectively stratified metabolic dysfunction-associated steatotic liver disease (MASLD) population outcomes. Graphical abstract for the identification, validation and translation of the flavoproteome as a target in MASLD.

## Results

### A digital network-based model nominates the FlavProt as a target in MASLD

MASLD affects diverse molecular networks across disease states and cell types, hampering the identification of therapeutic targets^2^. We developed *MetaLiver*, a multimodal network framework that integrates disease-, progression-, and cell type-specific information to reveal druggable targets. We first generated a disease-focused KG to contextualize candidate compounds within MASLD-relevant biology (MASLD-KG; **Fig. 2A**). The biomedical resource PrimeKG^10^ provides a foundation for network-based drug discovery, but lacks disease specificity and involves high computational costs. Therefore, we seeded the graph on the disease node for “metabolic dysfunction-associated steatohepatitis” (MASH), concentrating MASLD-biology (seed-only; **Fig. 2B**). Relevant terms including “lipid and fatty acid” or “insulin and glucose” showed an enrichment of 4.0-or 5.7-fold relative to the full graph, respectively (**Fig. 2B, Supp. Table 1**). Across all six families combined, the seed-only graph was the most MASLD-enriched of eleven subgraphs matched on size and degree, at 2.12-fold above its own matched null versus 1.91–2.07 for ten control diseases, including diabetes mellitus and coronary artery disease (**Fig. 2C**). The seed-only graph directly connected established MASLD clinical target genes, including *FGF21*^11^, *NR1H4*^12^, and *GLP1R*^4^. Strikingly, it required the incorporation of stratified MASLD patient data^13^ to connect *THRB*, the first and currently only liver-directed target^3^, which was not reachable from the disease node alone (**Fig. 2D**). This held true for the MASLD risk gene *MBOAT7*^14^ which, unlike other risk mutations such as *PNPLA3*^15^ or *TM6SF2*^16^, also required patient data incorporation (**Supp. Fig. 1A**). Further, patient-data integration expanded the graph 5.2-fold, from 1,063 to 5,534 genes, while retaining enrichment above all ten degree-matched random augmentations of the same scaffold (1.47 versus 1.32–1.43; **Supp. Fig. 1B**). Thus, clinical data extended the access to disease-relevant biology within the MASLD-KG beyond the seed-provided network. Importantly, despite increasing disease relevance, we reduced the overall computing time 18.8-fold (from 63.0 to 3.4 min) and the dominant betweenness-centrality computation 33.4-fold (from 63.0 to 1.9 min; exact Brandes, single-threaded), while retaining drug rankings of the full graph (Spearman rho 0.99) (**Fig. 2E, Supp. Fig. 1C**). To next identify compounds targeting key nodes in networks affected in MASLD-progression, we applied weighted gene co-expression network analysis (WGCNA^17^) to the patient data^13^ incorporated in the MASLD-KG. This strategy identified six co-expression modules significantly correlated with the disease progression marker fibrosis, including an extracellular matrix (ECM) gene module, validating this strategy (**Supp. Fig. 2A&B**). To derive target network nodes, we focused on the repressed ME8 module due to its enrichment in metabolic and mitochondrial processes, central in MASLD^18^ (**Supp. Fig. 2C**). As network analysis revealed a direct interaction for only a subset of the co-expressed genes (**Supp. Fig. 2D**), we used Dijkstra’s algorithm to identify the shortest paths between genes, capturing indirectly related nodes. This strategy identified a network comprising 7,545 proteins, with 70 key nodes that remained enriched for metabolic and mitochondrial processes (**Supp. Fig. 2E&F**). A network-based drug proximity model^19^ identified 132 significant FDA-approved compounds for targeting these key nodes (Z < –1.96; **Supp. Table 2**). Since the predicted compounds lacked cell type specificity, we intersected genes upregulated in MASLD livers^13^ with genes detected in a subset of MASLD-hepatocytes in mouse^20^ and human^21,22^ liver sc-/sn-RNA-seq data sets (genes without a mouse ortholog were assessed in human data only), yielding a 35-gene signature (**Supp. Fig. 3A-F**). From this signature, we built a protein-protein-interaction (PPI-) network using STRING and identified topological hubs by applying centrality metrics (**Supp. Fig. 3G**). Permutation testing refined this network to 21 high-confidence nodes that were both central and transcriptionally upregulated. These nodes distinguished steatotic liver patients from controls and were significantly enriched for the KEGG pathway “Non-alcoholic fatty liver disease” (**Supp. Fig. 3H-J**). Network-based drug proximity analysis identified 34 FDA-approved compounds with significant proximity to the 21-node disease module in the human interactome (**Supp. Table 3)**. We integrated these complementary network layers into the publicly accessible framework *MetaLiver*. *MetaLiver* prioritized FAD highly, followed by NADH and PLP, the functional cofactors of the vitamin B2-, B3-and B6-dependent networks (**Fig. 2F**). We chose the vitamin B2-dependent network, the FlavProt, for further investigation given that flavoproteins are known in energy homeostasis but are poorly defined in MASLD progression. Thus, *MetaLiver* offers the identification of tractable networks, e.g. the FlavProt in MASLD.

**Fig. 2:**
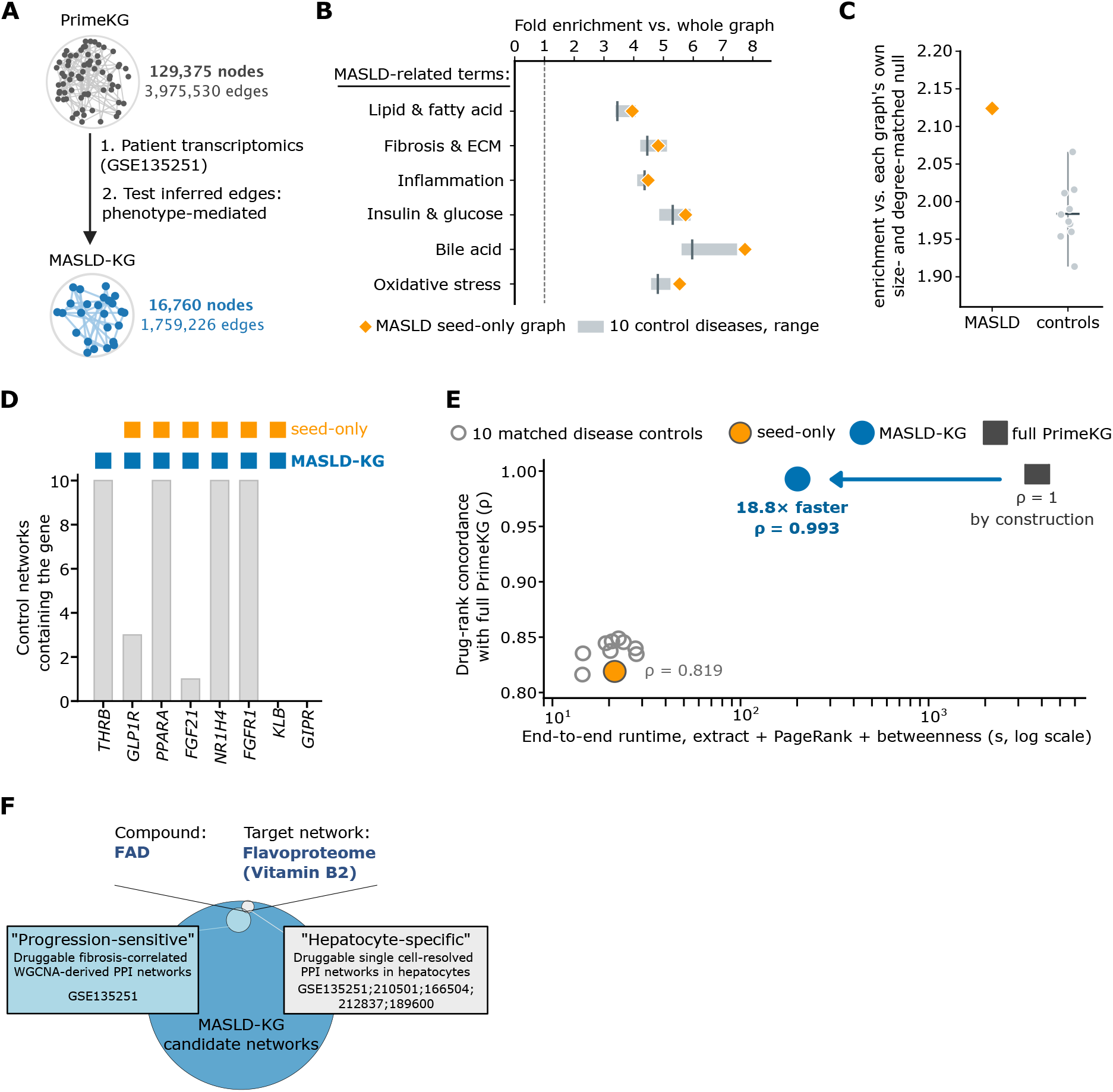
Knowledge graph-driven drug prioritization framework nominates the flavoproteome network as a target in MASLD. A) Strategy for generating the MASLD-specific knowledge graph (MASLD-KG). B) Fold change enrichment for MASLD-related pathway terms in the seed-only graph (grown from the “MASH” disease node alone; yellow diamonds), shown for each of the six MASLD pathway families (term counts in brackets). In grey, the range and median across ten control-disease subgraphs matched on size, degree profile and node class composition (n = 10). The dashed line marks fold enrichment = 1 (no enrichment above the whole-graph background) C) Enrichment across all six MASLD-relevant families shown in B) combined, expressed relative to a null matched on both gene-set size and log-degree profile, for the seed-only graph and the ten matched control diseases. D) Direct connection of known drug targets for treating MASLD in the seed only and clinical data-informed graph compared to other non-MASLD related KGs of same size (n=10) E) Performance of the MASLD-KG compared to PrimeKG, seed-only control and 10 other size-matched controls. F) Venn diagram of the drug prioritization strategy. Circle sizes reflect the number of candidate compounds.

### The FlavProt is dysregulated in MASLD models and can be therapeutically targeted

Based on *MetaLiveŕs* prediction, we hypothesized that progressing MASLD injury induces FlavProt dysregulation in hepatocytes. We therefore pursued an experimental model that progressively escalates metabolic stress in iPSC-and primary hepatocytes by integrating: 1) substrate overload through oleic and palmitic acid exposure (OA/PA), 2) extra-hepatic metabolic stressors recapitulating adipose (resistin^23^) and sarcopenic signaling (myostatin^24^) (Res/Myo), and 3) inflammatory stress and intestinal permeability by, e.g., indirectly co-culturing endotoxin-primed peripheral blood mononuclear cells (PBMCs) (**Fig. 3A**)^25,26^. Increasing injury progressively altered the FlavProt gene expression alongside established MASLD markers, including released aspartate aminotransferase (AST), TNFα and IL-1β, mitochondrial membrane potential (ΔΨm) depolarization, and steatosis (**Fig. 3A**). Res/Myo & PBMC-exposed Ci-Heps showed disturbances of FlavProt-dependent pathways including mitochondrial function that resembled transcriptomes of fibrosis stage 4 MASLD patients (**Supp. Fig. 4A**). Consistently, the culture supernatants contained clinical disease markers including DPP4^27^, LDHA and LDHB^28^ (**Supp. Fig. 4B, Supp. Table 4**). FlavProt-related disturbances, e.g., the increased expression of the vitamin B2 metabolizing enzyme *RFK* or ΔΨm depolarization, showed consistency across Ci-Heps from three individual iPSC donors (**Supp. Fig. 4C-E**). We next hypothesized that the FlavProt alterations resulted from the depletion of its required cofactor FAD, predicted by *MetaLiver* as a candidate compound. Surprisingly, FAD levels remained unchanged in Ci-Heps suggesting rather direct injury-induced reprogramming of the FlavProt (**Supp. Fig. 4F**). Strikingly, increasing intra-cellular FAD >1.5-fold through its supplementation mitigated ∼75% of the observed FlavProt gene expression changes (**Fig. 3B**, **Supp. Fig. 4F**). Beyond the direct FlavProt effects, FAD supplementation (FAD^+^) reversed injury-induced transcriptomic changes in Ci-Heps including ECM-interaction or suppressed metabolic pathways (**Fig. 3C, Supp. Fig. 5A&B, Supp. Table 5**). To functionally test the therapeutic potential of FlavProt-targeting, we evaluated established hallmarks of MASLD. FAD^+^ reduced steatosis in Ci-Heps, most robustly from 50 µM onward compared to baseline FAD conditions (**Supp. Fig. 5C&D**). We validated the anti-steatotic effects in Ci-Heps across three individual primary human hepatocyte donors (**Fig. 3D**). Besides total steatosis, FAD^+^ also restored beneficial endogenous saturated-to-unsaturated and omega-3-to-omega-6 fatty acid ratios^29^ in Ci-Heps (**Fig. 3E, Supp. Fig. 5E**). Mitochondrial dysfunction is another key hallmark of MASLD contributing to disease progression^30^. FAD^+^ significantly increased ΔΨm in primary human Ci-Heps (**Fig. 3F**). It specifically induced the activity of mitochondrial complex II, suppressed in MASLD patients^30^ (**Fig. 3G**), potentially mediating this effect. For proof-of-concept expansion of the therapeutic scope beyond direct FAD^+^, we harnessed *MetaLiver* to derive pharmacological compounds targeting similar network nodes. We prioritized those enriched in hepatocytes under disease-like conditions, curating drug–target annotations for compounds targeting both the FAD module and its proximal, hepatocyte-relevant neighborhood (**Supp. Table 6**). Of the five experimentally tested candidates, norleucine, 3-Pyridin-4-yl-1H-indazol-5-ylamine dihydrochloride (3P), and aspirin fully restored mitochondrial spare respiratory capacity in Ci-Heps similarly to FAD^+^ (**Fig. 3H, Supp. Fig. 5F**). Notably, 3P increased the expression of the vitamin B2 homeostasis genes *SLC52A2*, *RFK* and significantly *FLAD1*, potentially mediating this effect (**Supp. Fig. 5G**). Ultimately, we tested the impact of FAD^+^ on the immuno-metabolic coupling relevant in MASLD^31^ exposing human PBMCs to the supernatant of the PBMC-containing Ci-Hep model (18 h) (**Supp. Fig. 6**). The supernatant robustly induced CD40 and CD80 on CD14^+^ monocytes, which are crucial co-stimulatory proteins for T-cell activation (**Supp. Fig. 6A&B**). Consistently, PD-1 and CD25, which transiently accompany early T-cell activation^32,33^, showed an increased surface expression on CD4^+^ and CD8^+^ T-cells (**Supp. Fig. 6C-E**). FAD^+^ significantly inhibited this immune cell activation, which persisted during LPS-mediated activation. Together, these analyses demonstrate that metabolic stress, rather than cofactor deprivation, induces FlavProt dysregulation, thereby promoting cellular injury that can be mitigated by cofactor replenishment or network-modifying drug compounds.

**Fig. 3:**
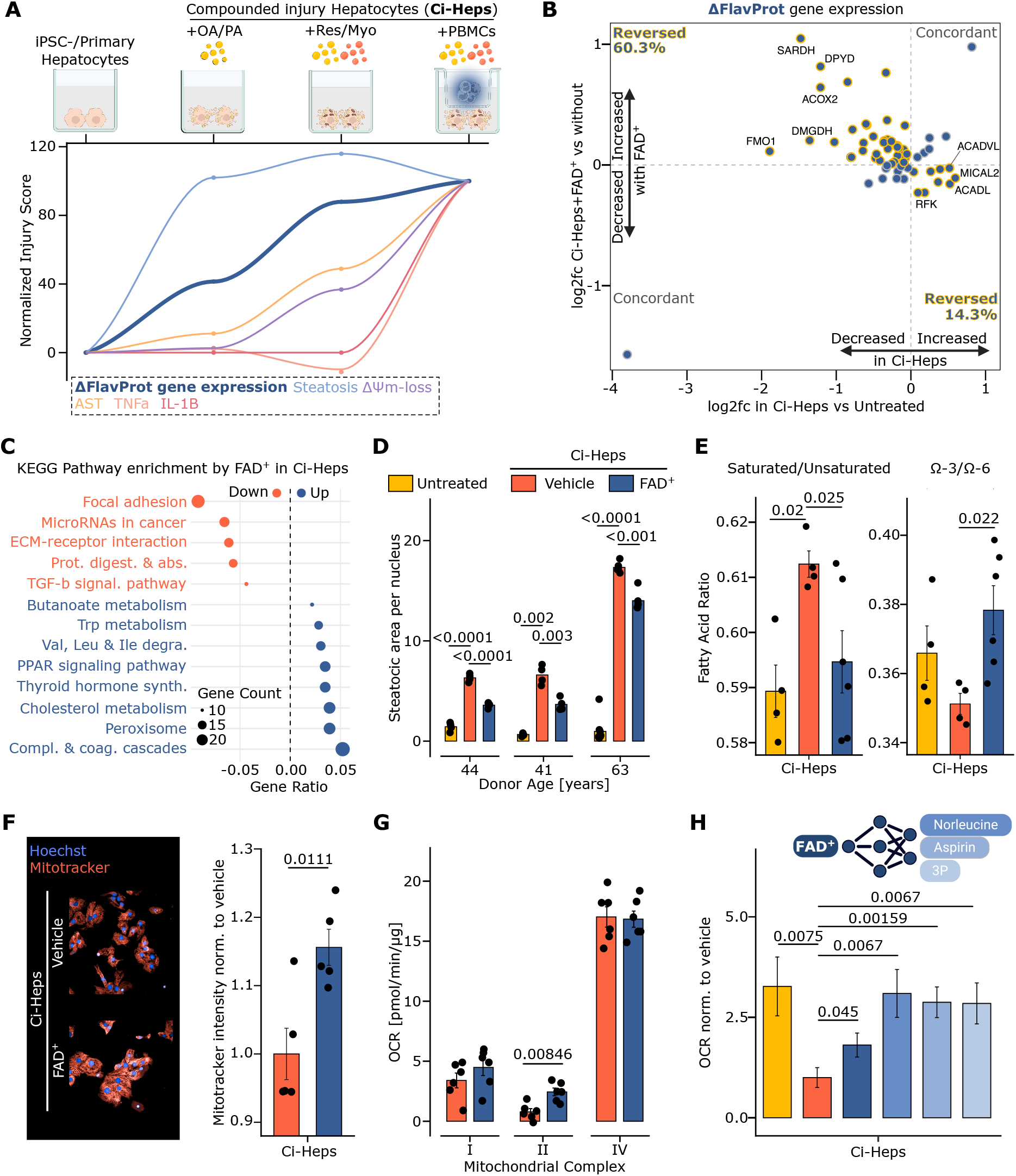
The flavoproteome (FlavProt) is functionally perturbed in preclinical MASLD models and its cofactor replenishment shows mitigation of metabolic injury. A) Gene expression changes of the FlavProt network and hallmark MASLD markers in a compounded metabolic stress model using induced pluripotent stem cell (iPSC-) derived hepatocytes. All parameters were standardized, ranging from untreated (0%) to the most complex disease condition including all disease triggers (100%). Pooled FlavProt gene expression changes were assessed analyzing RNA seq data for all genes from the human FlavProt across disease conditions. TNFa, IL1B and AST protein data are from supernatant. Steatosis and mitochondrial membrane potential (ΔΨm) was assessed using live dye-based assays. Each parameter was assessed in n ≥ 3 independent samples. Oleic and palmitic acid = OA/PA; Resistin and myostatin = Res/Myo; Peripheral blood mononuclear cells = PBMCs B) Relative gene expression changes for all genes from the human FlavProt in iPSC-Ci-Heps exposed to injury with or without 50 µM FAD (FAD^+^). The x-axis shows relative expression changes in iPSC-Ci-Heps (OA/PA, Res/Myo, PBMCs) compared to untreated control. The y-axis shows relative expression changes in iPSC-Ci-Heps (OA/PA,Res/Myo,PBMCs) compared to iPSC-Ci-Heps with FAD^+^. Reverse directionality is indicated by yellow outlines. Changes are shown for n ≥ 3 independent samples. C) Top significant KEGG pathways enriched in iPSC-Ci-Heps exposed to OA/PA, Res/Myo and PBMCs treated with FAD^+^ relative to the vehicle control. Changes are shown for n ≥ 3 independent samples. Prot. digest. & abs. = Protein digestion & absorption; TGF-b signal. pathway = TGF-b signaling pathway; Val, Leu & Ile degra. = Val, Leu & Ile degradation; Compl. & coag. cascades = Complement & coagulation cascades D) Quantification of steatotic area per nucleus in primary human hepatocyte (PHH-) Ci-Heps derived from different donors. Ci-Hep injury refers to OA/PA, Res/Myo and the pro-inflammatory cytokines IL1B and TNFa. Each dot represents an independent sample. P-values corrected using the Benjamini-Hochberg method after applying multiple t-tests are shown. E) Quantification of the saturated over non-saturated and omega-3 over omega-6 fatty acid ratios normalized to protein concentration in iPSC-Ci-Heps. Ci-Hep injury refers to Res/Myo and PBMC co-culture. Each dot represents an independent sample. P-values corrected using the Benjamini-Hochberg method after applying multiple t-tests are shown if significant. F) Representative images and quantification of ΔΨm in PHH-Ci-Heps. Ci-Hep injury refers to OA/PA, Res/Myo and the pro-inflammatory cytokines IL1B and TNFa. ΔΨm was normalized to the vehicle control. Each dot represents an independent sample. P-value after applying a t-test is shown. **G)** Mitochondrial complex activity (I, II and IV) in primary hepatocyte Ci-Heps. Ci-Heps injury refers to OA/PA and Res/Myo. Each dot represents an independent sample. P-values corrected using the Benjamini-Hochberg method after applying multiple t-tests are shown if significant. **H)** Testing of mitochondrial spare respiratory capacity for translational compounds in primary hepatocyte Ci-Heps. Mitochondrial spare respiratory capacity was assessed using the Seahorse XF Cell Mito Stress Test. Norleucine (Nor) was used in a 1 mM, Aspirin (Asp) in 5 µM and 3-Pyridin-4-yl-1H-indazol-5-ylamine dihydrochloride (3P) in 1.5 µM concentration. For each group, at least 12 independent samples were included. Ci-Hep injury refers to OA/PA and Res/Myo. P-values corrected using the Benjamini-Hochberg method after using multiple t-tests are shown.

### A FlavProt signature stratifies MASLD progression identifying AIFM2 as a critical bottleneck

Having validated the therapeutic tractability of FlavProt dysregulation in pre-clinical models, we next asked whether this axis shows clinical relevance in MASLD patients. We therefore analyzed the clinical MASLD cohort SteatoSITE^34^, which we intentionally kept as validation cohort, not included in the *MetaLiver* prediction (**Fig. 4A**). SteatoSITE comprises matched liver histopathology, comprehensive clinical annotation, and bulk transcriptomic profiles from MASLD patients^34^. Using LASSO-penalized Cox regression in 446 patients including all FlavProt genes, we identified a 19-gene module significantly stratifying patients with hepatic decompensation risk. Strikingly, the resulting FlavProt score showed a similar performance strength as the clinically established fibrosis marker FIB-4, displaying a 1.56-fold increase in hazard ratio (**Fig. 4B&C**). Further, based on hazard ratio, the FlavProt score showed the best performance over other tested gene networks, including ECM and niacin (vitamin B3-) dependent genes, given their recently described crucial role in liver regeneration^35^ (**Fig. 4C**). The FlavProt score comprises decreased and increased gene expression levels indicating both loss-of-function and compensatory transcriptional responses (**Fig. 4D**). Besides genes directly connected to FAD homeostasis, including reduced *RFK* expression or ectopic expression of the intestinal vitamin B2 transporter *SLC52A3*^36^, decreased *AIFM2* expression was one associated risk feature. Recent work identified the oxidoreductase AIFM2 as a ferroptosis inhibitor in cancer cells^37^, also now known as FSP1, without a functionally defined role in MASLD. We therefore assessed the impact of pharmacological AIFM2-inhibition using iFSP1, a small molecule inhibitor that blocks the ubiquinone, not the FAD, binding pocket^38^ of AIFM2, in metabolically stressed Ci-Heps with or without FAD^+^ (**Fig. 4E**). AIFM2-inhibition sensitized Ci-Heps to ferroptotic stress indicated by significantly decreased cell viability (**Fig. 4F**). Conversely, FAD^+^ abolished this effect (**Fig. 4F**). Since AIFM2 controls lipid peroxidation^37^ thereby inhibiting oxidative stress, we assessed reactive oxygen species (ROS) levels and steatosis. FAD^+^ significantly reduced ROS levels independent of AIFM2-activity, potentially via increased glutathione regeneration augmenting GPX4-dependent cell protective pathways^39^ (**Fig. 4G, Supp. Fig. 7A**). In contrast, inhibiting AIFM2 attenuated the anti-steatotic effects of FAD^+^, implicating uncoupled mechanisms for hepatocyte death-protection and steatosis resolution (**Fig. 4H**). This data nominates AIFM2 as a mechanistic node for the FAD^+^-induced reduction in lipid accumulation. Together, these findings establish the clinical relevance of FlavProt dysregulation and nominate critical bottleneck genes for MASLD exacerbation. Our data propose reduced activity of the oxidoreductase AIFM2 as a critical risk factor for MASLD-associated hepatic decompensation, mediated via an increased vulnerability for hepatocyte death. Importantly, FAD^+^ protected hepatocytes under AIFM2-deficiency underscoring the potential of the vitamin B2-dependent therapeutic axis.

**Fig. 4:**
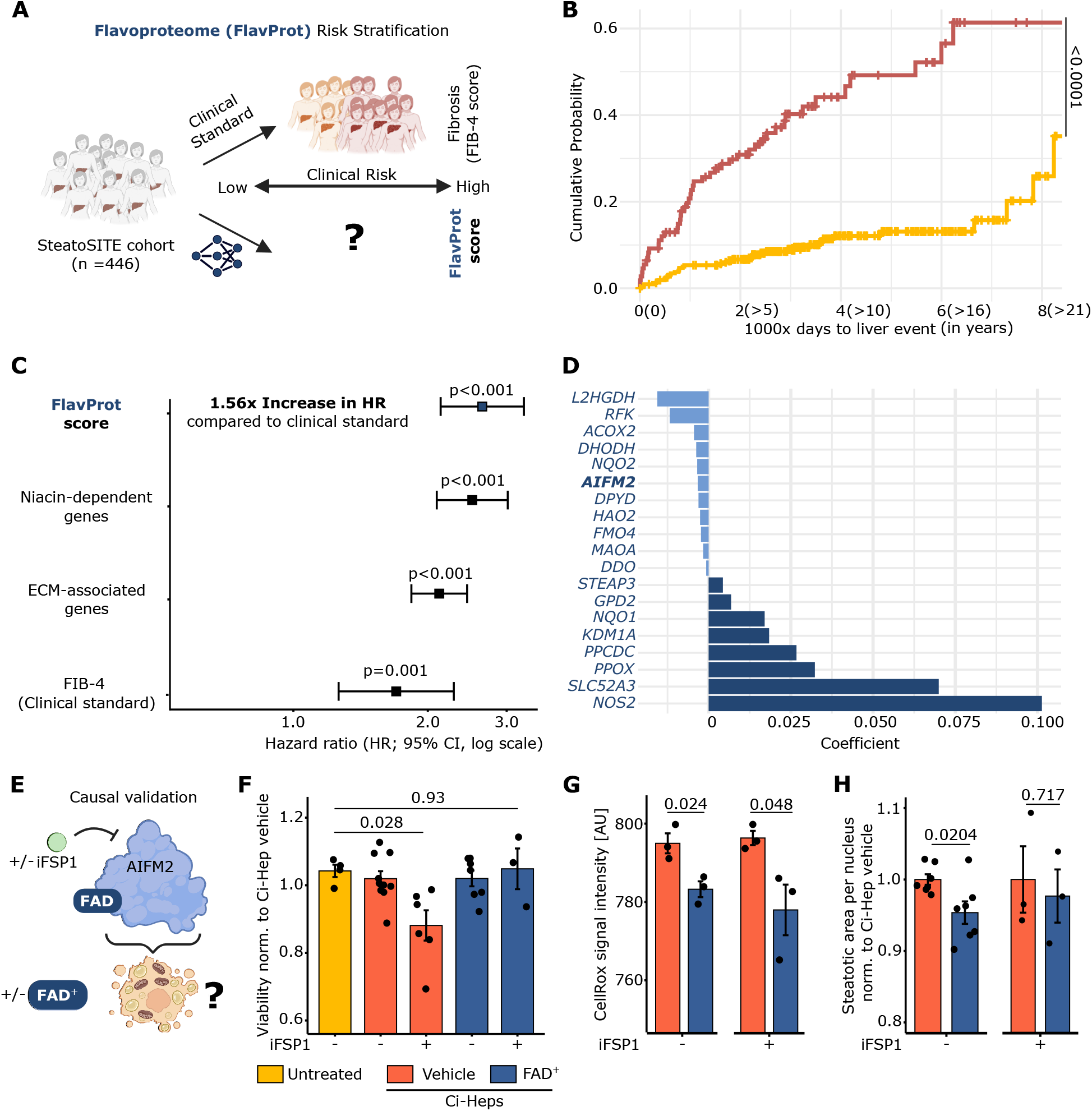
FlavProt expression score stratifies MASLD outcome and identifies AIFM2 deficit mediated hepatocyte injury. A) Graphical summary of FlavProt score identification B) FlavProt-based classification performance for hepatic event probability using LASSO-penalized Cox regression. Significance was assessed using a log-rank test. Red = high risk patient. Yellow = low risk patient. C) Discriminative performance of the FlavProt score compared to other gene signatures, including extra cellular matrix (ECM) genes and niacin (VitB3-) dependent genes, and the clinical fibrosis marker FIB 4. D) Gene expression coefficients for the FlavProt score nominating clinical bottleneck genes. AIFM2 is highlighted. E) Graphical depiction of the experimental set up to test AIFM2-inhibition in hepatocytes undergoing MASLD injury. F) Viability in induced pluripotent stem cell (iPSC-) derived hepatocytes assessed via the PrestoBlue assay normalized to the vehicle control without iFSP1. Compounded injury in hepatocytes (Ci-Heps) refers to oleic and palmitic acid (OA/PA) exposure and peripheral blood mononuclear cell co-culture (PBMCs). Each dot represents an individual sample G) Quantification of reactive oxygen species (ROS) intensity in iPSC-Ci-Heps (OA/PA&PBMCs). Each dot represents and individual sample. P-values corrected using the Benjamini-Hochberg method after applying multiple t tests are shown. H) Quantification of steatotic area in iPSC-Ci-Heps (OA/PA&PBMCs) normalized to nucleus count and the vehicle control. Each dot represents and individual sample. P-values corrected using the Benjamini-Hochberg method after applying multiple t-tests are shown.

## Discussion

After several Phase 3 clinical trials, only Resmetirom and Semaglutide received FDA approval for treating MASLD. Still, only a subset of patients shows a response to both drugs, emphasizing the need for additional treatment strategies^3,4^. Conventional drug development, however, requires time and costs, which can be reduced by digital prioritization^41^. Here we built a large *in silico* model fed with patient data, interrogated against a stratified MASLD cohort, and then functionally checked for mechanistic bottleneck nodes *in vitro*. Specifically, we introduce *MetaLiver,* an open-access platform to identify active biological networks critical for drug development and repurposing. *MetaLiver* comprises a KG specific for MASLD, complemented with patient fibrosis-correlated and hepatocyte-specific network analysis. This framework identified the FlavProt as a target network. We validated its functional relevance using iPSC-derived and primary hepatocytes and immune cells. Using a large-scale MASLD patient cohort, we link FlavProt dysregulation to defined clinical outcomes and nominate candidate bottleneck genes. Of these, we functionally establish *AIFM2* as a key FlavProt node whose inhibition exacerbates hepatocyte death in MASLD, mitigatable via increased vitamin B2 fueling.

ML-driven strategies increasingly transform drug development and repurposing^5^. For example, a recent study applied large language model (LLM-) assisted drug repurposing to derive anti-fibrotic drugs, validated in liver organoid models^42^. We here utilize a complementary strategy for drug prioritization based on biomedical KGs. We overcame existing limitations of PrimeKG by expanding relevant disease nodes and integrating patient data, generating a MASLD-specific KG. This strategy increased biological specificity, with the curated MASLD neighborhood being the most disease-enriched of eleven size-and degree-matched disease subgraphs and connected clinical target and risk genes not associated before. While disease-specific KGs exist, e.g., for Alzheimeŕs^43^, this is, to our knowledge, the first MASLD-focused KG allowing complex translational associations. Recent strategies also combine LLM- and KG-based approaches for cancer drug repurposing, which could be iteratively implemented in future MASLD drug prioritization frameworks^44,45^. By applying additional network-based algorithms, we identified progression-sensitive metabolic- and hepatocyte-specific druggable therapeutic nodes. Together, we develop a multimodal and iterative strategy for druggable network prioritization, thereby substantially advancing earlier frameworks^46,47^. We then validate one of *MetaLiver*’s predictions, the FlavProt, as a critical and therapeutically targetable network. For experimental testing, we chose non-transformed hepatocyte systems since malignant models exhibit drastic metabolic rewiring^48^. We demonstrate FlavProt dysregulation by MASLD stressors, including resistin and myostatin, which thus far have not been described in that context. Thus, supporting a new angle to their debated role in human MASLD, advancing prior reports from mice and carcinoma cell lines^23,49–52^. Importantly, we show that metabolic stress causes the FlavProt changes despite basal cofactor availability, distinguishing our findings from previous studies that induced vitamin B2 deficiency in rodents^53,54^, zebrafish^55^, and HepG2 cell^53^ models. We intentionally chose FAD supplementation as the primary source for vitamin B2, given its higher aqueous solubility^56^ and cell-membrane-localized conversion to riboflavin by ecto-5’ nucleotidase CD73 and alkaline phosphatase to enter the cells via SLC52A transporters^57^. Despite early reports of cofactor reduction in end-stage liver disease^58^, this is, to our knowledge, the first study to demonstrate FlavProt dysregulation and its therapeutic effects in human MASLD. Ultimately, we establish the clinical relevance of the FlavProt using a retrospective, stratified MASLD patient cohort. We identify a polygenic FlavProt expression module identifying patients at high risk for hepatic decompensation. Given the growing cost-effectiveness of transcriptomic profiling, such strategies could help inform personalized clinical decision-making, already exemplified in clinical oncology^59–61^. In the future, prospective, independent multicenter validation may support the FlavProt score to be widely adopted for MASLD risk stratification. The FlavProt score nominated reduced *AIFM2* expression as a key mediator for increasing hepatic decompensation risk. Recent studies identified AIFM2, now also recognized as FSP1, in cancer cell lines as a vitamin B2-dependent ferroptosis inhibitor regulating survival^62,63^. Vitamin B2-derived FAD is thereby crucial to stabilize AIFM2 and to enable its enzymatic activity. Interestingly, AIFM2 abundance increased after knocking out the E3 ubiquitin-protein ligase *RNF126*^63^, which in our data is a key node in the hepatocyte-specific MASLD-network (**Supp. Fig. 3H**). Thus suggesting that RNF126 may enhance AIFM2 degradation in hepatocytes, increasing their MASLD vulnerability. However, future RNF126 perturbations are needed to validate this hypothesis, as contrasting roles for this protein have been described, including mediating the re-localization of AIFM2 and thereby causing an increased ferroptosis resistance in medulloblastoma^64^. We here extend the current knowledge about AIFM2 beyond cancer cell biology, showing its crucial role in protecting non-transformed hepatocytes under MASLD-associated stress. Importantly, FAD^+^ mitigated the increased susceptibility to cell death under AIFM2 inhibition, suggesting independent buffering effects, potentially by reducing the overall oxidative load via improved glutathione regeneration for the auxiliary GPX4 pathway^39^. Notably, iFSP1 has been described to bind the ubiquinone rather than the FAD binding pocket^38^, supporting that the observed rescue effects are mediated via alternative molecular pathways rather than competition-based AIFM2 re-activation. Mechanistically, FAD^+^’s anti-steatotic effects required intact AIFM2 activity, nominating it as a functional FlavProt effector for this process. However, the sustained steatosis did not abolish the hepatocyte-protective effects, implicating it rather as a benign cellular response than an injury-driver. In the future, genetic knock-out or -down strategies will further validate our findings and help to decipher the underlying molecular mechanisms of how AIFM2 mediates the anti-steatotic effects, especially given its lipid droplet localization^65^ and role in controlling lipid droplet quality^66^. Together, these data position AIFM2 as a hepatocyte-protective factor implicated in liver decompensation, whose critical loss-of-function phenotype can be compensated by activating vitamin B2-dependent signaling.

Besides vitamin B2-, *MetaLiver* also predicted B3- and B6-dependent networks as therapeutic targets in MASLD. This prediction aligns with recent studies showing beneficial B3-mediated effects on liver regeneration^35^ or treating MASLD^67^, or B6 in a non-placebo-controlled single-center study reducing hepatic steatosis in MASLD patients^68^. Functional gain- or loss-of-function nutrigenomic approaches, as shown for cancer cell proliferation^69^, will further help decipher the specific vitamin impact on complex metabolic diseases including MASLD. Notably, high-dose vitamin B2 is clinically applied for rare diseases^70^ and migraine prevention^71^, facilitating its evaluation in treating MASLD as done for vitamin E^72^. However, whether existing dosing regimens will reach therapeutic concentrations in the liver remains to be evaluated.

In summary, we establish the multimodal framework *MetaLiver* to guide and accelerate target network identification for MASLD. We exemplify its iterative optimization and hypothesis validation using complex human cell models and clinical data, thereby identifying the FlavProt as a targetable network. Extending this framework toward synergistic multi-pathway targeting or patient-specific predictions could further advance the dissection of complex diseases and enable more precise therapeutic interventions.

## Materials and methods

### Study overview and ethics statement

We developed a ML-based framework to identify druggable networks in MASLD and validate one predicted network using orthogonal experimental approaches and clinical data.

We applied three complementary network-based approaches capturing different public data sources and modalities, to emphasize the multi-facetted pathophysiology of MASLD:

(1) Generation of a MASLD-specific KG to uncover systems-level relationships linking drugs and hepatic phenotypes

(2) fibrosis-resolved WGCNA to capture disease progression signals

(3) PPI-based network proximity analysis grounded in single cell transcriptomics specific for hepatocytes. This pipeline enabled prioritization of networks and bioactive molecules with translational potential for intercepting MASLD. We focused on the FlavProt for validation given its ill-defined role in MASLD. We tested network manipulation using hepatocyte-centric model systems, in which FAD supplementation served as primary intervention. We cross-validate findings using iPSC-derived and primary human hepatocytes, mouse hepatocytes, and primary human immune cells. To facilitate clinical translation, we further predicted pharmacological compounds targeting similar network nodes as FAD and validated their bioactivity *in vitro*. Ultimately, we use a large independent cohort of MASLD patients (n=446) to connect FlavProt network disturbances with clinical outcomes. We thereby derive candidate FlavProt bottleneck genes, of which we validate one candidate experimentally for its relevance and mechanistic influence. All animal experiments complied with ethical regulations and were approved by the Committee on Research Animal Care of the Charité Universitätsmedizin Berlin, Berlin, Germany (License number: G0174/20). Human cell isolation was done in accordance with the amended Declaration of Helsinki. Specifically, primary hepatocyte isolation was done under informed consent and the ethics statement EA1/257/14. Blood isolation was done in accordance with ethical standards of the Charité Universitätsmedizin Berlin under the ethical approval EA2_065_21. SteatoSITE data were derived from anonymized tissue samples approved for use under the NHS Research Scotland Biorepository Network (SR1032) and the unified pan-Scotland governance framework for unconsented data inclusion (West of Scotland REC 4: 20/WS/0002; PBPP: 1819-0091)^34^

### Statistical testing

We performed data visualization and statistical significance assessment using primarily RStudio or additional software as indicated. If not stated otherwise, we applied two-sided, multiple t-test with p-values that we corrected using the Benjamini-Hochberg method. The n-numbers are described in each figure legend. Unless stated otherwise, all bar plots show mean and standard deviation.

## Data availability

The main data supporting the results in this study are available within the article and its supplementary information. Raw data of the *in vitro* transcriptomics will be provided to the reviewers if requested and uploaded to a public repository upon publication. The SteatoSITE transcriptomic data are deposited in the European Nucleotide Archive (https://www.ebi.ac.uk/ena; study accession number: PRJEB58625). Gene expression data are also freely available for user-friendly interactive browsing online at https://shiny.igc.ed.ac.uk/SteatoSITE_gene_explorer/. Full data access can be requested at https://regeneration-repair.ed.ac.uk/research/research-resources/steatosite. Mass Spec raw data can be found as indicated. Further, all data sets incorporate in *MetaLiver* can be interactively explored at https://gehadyoussef.github.io/meta-liver/ The code for the described analyses can be found here https://github.com/Namshik-Han-Lab/MASH

## List of supplementary materials

Supplementary materials and methods including supplementary references and supplementary figures Supplementary tables S1 to S6

## Acknowledgments

We thank the members of the BIH Core Unit for Pluripotent Stem Cells and Organoids, specifically Valeria Fernandez Vallone, Janine Cernoch, and Claudia Schaar, for their technical support. We also thank Janine Altmüller and Tatiana Borodina of the MDC/BIH Genomics Platform, Berlin, for technical support. Additionally, we would like to thank the HPC for Research/Clinic cluster at the Berlin Institute of Health for providing data storage. The authors would like to acknowledge Novogene UK (Munich, Germany) for providing bulk RNA sequencing and bioinformatics analysis services. We are also grateful for the support provided by all the current and former members of the Rezvani Lab, specifically Natalia Martagón Calderón and Reinhild Dünnebacke. We would also like to thank the following people for sharing their expertise: Marième Gueye, Dennis Weschka Pilic, Amr Al Shebel, Tilman Breiderhoff, Kerstin Sommer, Sandra Gerstenberg, Verena Klämbt, Marah Alhabahbeh, Adrien Guillot, Hanyang Liu, Tian Lan and Guo Yin. Further, we want to thank Aras Mattis and lab members from the University of California, San Francisco, for discussions, and Leon Louis Seifert for his critical and constructive assessment of this work. Also, we would like to thank David Meierhofer. Furthermore, we would like to extend our deepest gratitude to the individuals who have donated materials to this biomedical research.

## Author contributions

M.R. and Na.H. conceived the study. M.R., J.W., Na.H., F.B., G.Y. and A.C. designed and supervised the study. K.L. and H.S. contributed to protocol establishment. Ni.H., I.S. and P.T. provided and isolated clinical specimens. P.K. and C.E. contributed SeaHorse analysis. S.Q. and M.M. contributed to *in vitro* Ci-Hep assays. M.Ro. and L.V. contributed to FoP-Hep generation. M.K. and S.E. contributed the Mass spec analysis. M.Rot. contributed the Lipidomics analysis. I.L. and L.H. contributed with the full spectrum flow cytometry analysis. T.K. and J.F. analyzed the SteatoSITE co-hort. B.C., C.H., T.T. F.T., P.R. and P.B. participated in discussions and provided intellectual input. J.W., M.R., Na.H., F.B., G.Y. and A.C. wrote the paper.

