## Supplementary Information for "Network modeling identifies vitamin B2 control of AIFM2 deficit-driven hepatocyte death in MASLD"

### Supplementary Figures

#### Supplementary Figure 1:

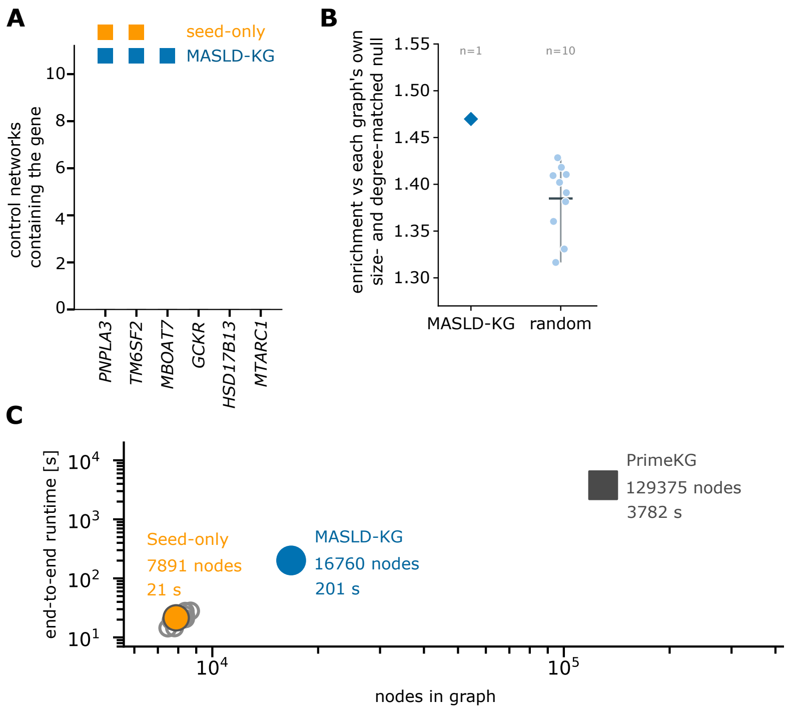


**Supp. Fig. 1: A disease specific knowledge graph increases relevance for metabolic dysfunction-associated steatotic liver disease (MASLD) while reducing computing costs. A)** Direct connection of known risk genes associated with MASLD in the seed-only and clinical data-informed graph compared to other non-MASLD related KGs of same size (n=10) **B)** The same measure as in **Fig. 2C** for the MASLD-KG against ten degree-matched random augmentations of the same scaffold. **C)** Performance of the MASLD-KG compared to PrimeKG, seed-only control and 10 other size-matched controls.

Supplementary Figure 2:

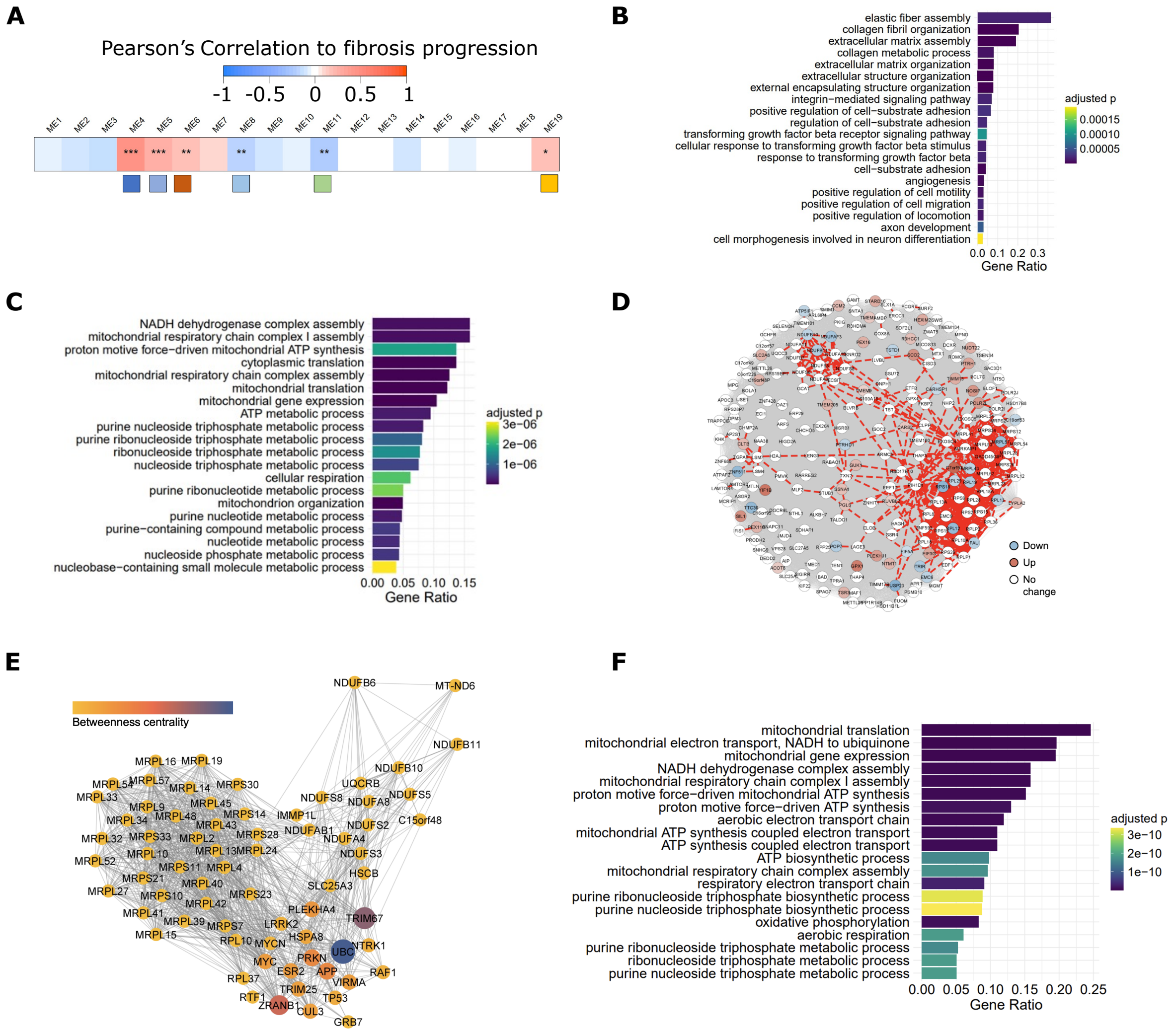

**Supp. Fig. 2: Fibrosis-correlated co-expression network analysis identifies therapeutic candidates for targeting metabolic and mitochondrial dysregulation in MASLD.** Weighted gene co-expression network analysis (WGCNA) was applied to transcriptomic profiles from the Govaere et al. cohort to identify gene modules associated with fibrosis progression in MASLD and nominate drug targets. **A)** Correlation heatmap between module eigengene expression and fibrosis stage. Each row represents a gene module (ME1–ME19), with Pearson correlation coefficients shown in the heatmap. Asterisks indicate statistical significance (* $p < 0.05$, ** $p < 0.01$, *** $p < 0.001$). Enriched biological processes for significantly correlated modules are indicated with colored labels (Dark blue = ECM organisation; blue = Proteosomal degration; orange = mRNA transport; light blue = Mitochondrial gene expression; green = Vesicle transport; yellow = RNA splicing). **B)** Top 20 enriched Gene Ontology (GO) terms for WGCNA module ME4, which was positively correlated with fibrosis severity. Terms include extracellular matrix organization, integrin signaling, and cellular migration. Bars are ranked by gene ratio and colored by adjusted p-value. **C)** Top 20 enriched GO terms for module ME8, ranked by adjusted p-value. ME8 was enriched for mitochondrial and translational processes, including NADH dehydrogenase complex assembly, oxidative phosphorylation, and ATP metabolic processes. Gene ratio represents the proportion of ME8 genes annotated to each GO term. **D)** Network visualization of ME8 showing both co-expression (grey edges) and physical protein–protein interactions (red edges). Nodes represent ME8 genes, colored by differential expression: red = upregulated, blue = downregulated, white = unchanged (relative to control). The network shows that only a subset of co-expressed genes physically interact, suggesting a layered structure of molecular coordination. **E)** PPI subnetwork constructed from ME8 hub genes after network expansion. Nodes are scaled by degree and colored by betweenness centrality. This subnetwork includes direct and indirect interactors connecting ME8 genes, used for drug proximity analysis. **F)** GO term enrichment for the 70 key nodes extracted from the ME8 expanded network. Terms include mitochondrial translation, respiratory chain assembly, ATP biosynthesis, and nucleotide metabolic processes. Enrichment remains highly specific to mitochondrial function, indicating that key biological themes are preserved after network expansion.

Supplementary Figure 3:

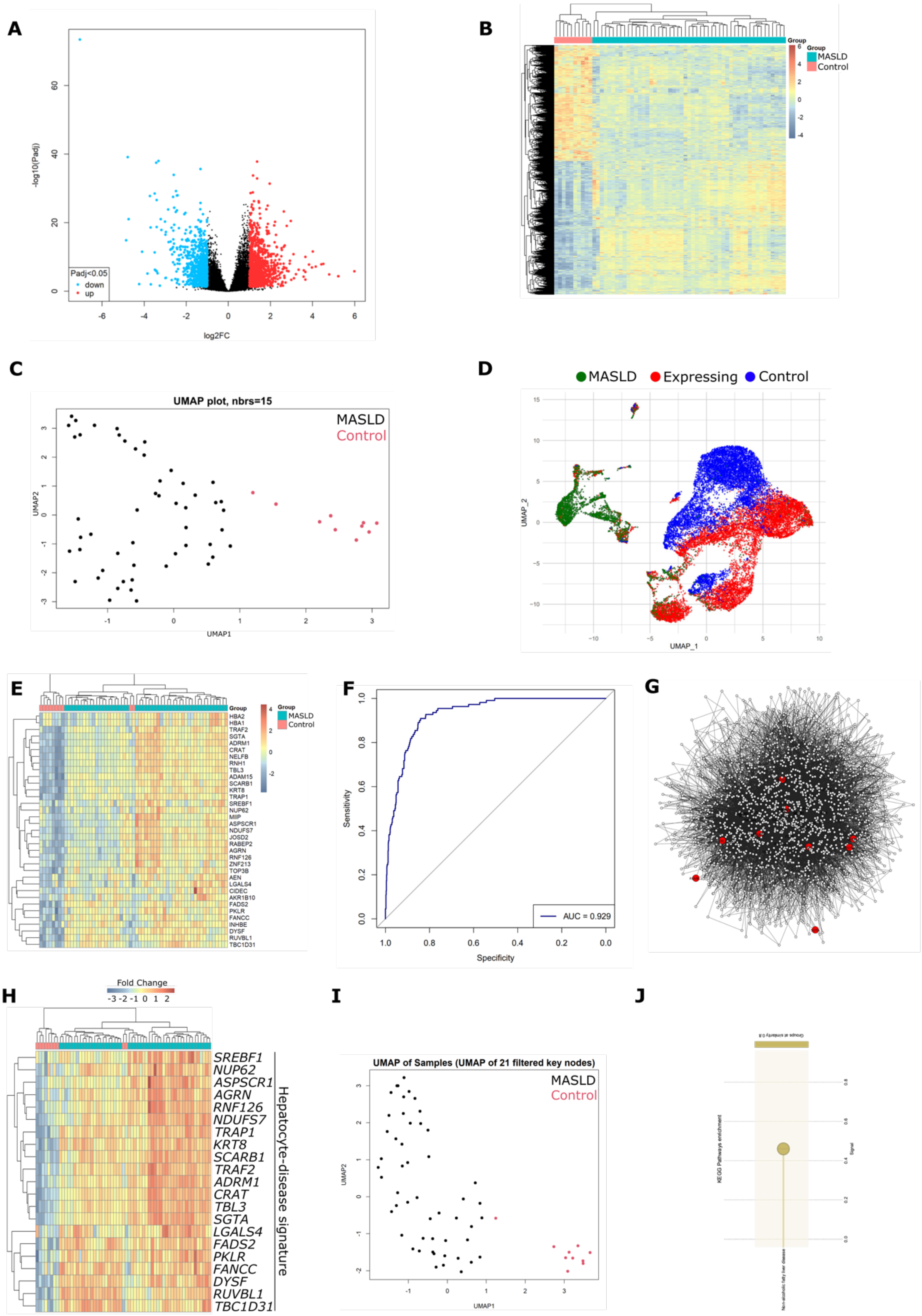

**Supp. Fig. 3: Identification of hepatocyte-specific MASLD signatures and drug targets via integrated transcriptomic and network-based analysis.** A multi-step pipeline was applied to derive a hepatocyte-specific disease module and prioritize drug candidates based on network proximity. **A)** Volcano plot showing differential expression analysis between early MASLD and control samples. Significantly upregulated (red) and downregulated (blue) genes are highlighted (adjusted $p < 0.05$), with log2 fold change on the x-axis and –log10 p-value on the y-axis. **B)** Heatmap showing the expression of 1,927 differentially expressed genes (DEGs) between early MASLD and control samples from human liver bulk RNA-seq data (GSE135251). Columns represent individual samples grouped by condition (early MASLD in red, control in turquoise), and rows represent DEGs scaled by row Z-score. Genes are hierarchically clustered. **C)** UMAP projection of bulk RNA-seq samples (GSE135251) based on global gene expression profiles. Samples cluster by condition, with early MASLD (red) clearly separated from control (black), indicating robust transcriptomic differences. **D)** UMAP projection of a representative liver single-cell RNA-seq dataset showing major cell populations colored by group: early MASLD (green), control (blue), and early MASLD signature-expressing cells (red overlay). Early MASLD-specific DEGs are predominantly expressed in hepatocyte clusters. **E)** Heatmap of hepatocyte-specific genes retained after cross-validation with four single-cell and single-nucleus RNA-seq datasets (mouse and human). This refined 35-gene early MASLD signature displays strong separation between early MASLD and control samples, confirming disease relevance and cell type specificity. **F)** Receiver operating characteristic (ROC) curve demonstrating the classification performance of the refined hepatocyte-specific MASLD signature. The area under the curve (AUC) is 0.929, indicating strong discriminative power. **G)** PPI network constructed from the refined early MASLD signature using STRING. Nodes represent proteins; edges indicate interactions. Red nodes highlight genes identified as network hubs by topological metrics (degree, closeness, betweenness, eigenvector centrality), filtered using permutation-based significance testing (N = 1,000). **H)** Expression heatmap of the 21 hub genes in the bulk RNA-seq dataset. Z-score-normalized rows illustrate consistent upregulation across early MASLD samples (red) and lower expression in controls (blue), further validating their disease association. **I)** UMAP embedding of bulk RNA-seq samples based on the expression of 21 key network nodes identified from the PPI analysis. The filtered signature maintains separation between early MASLD (red) and control (black) groups. **J)** KEGG pathway enrichment analysis of the 21 key PPI nodes. The only significantly enriched term was “Non-alcoholic fatty liver disease,” supporting the specificity of the derived subnetwork.

### Supplementary Figure 4:

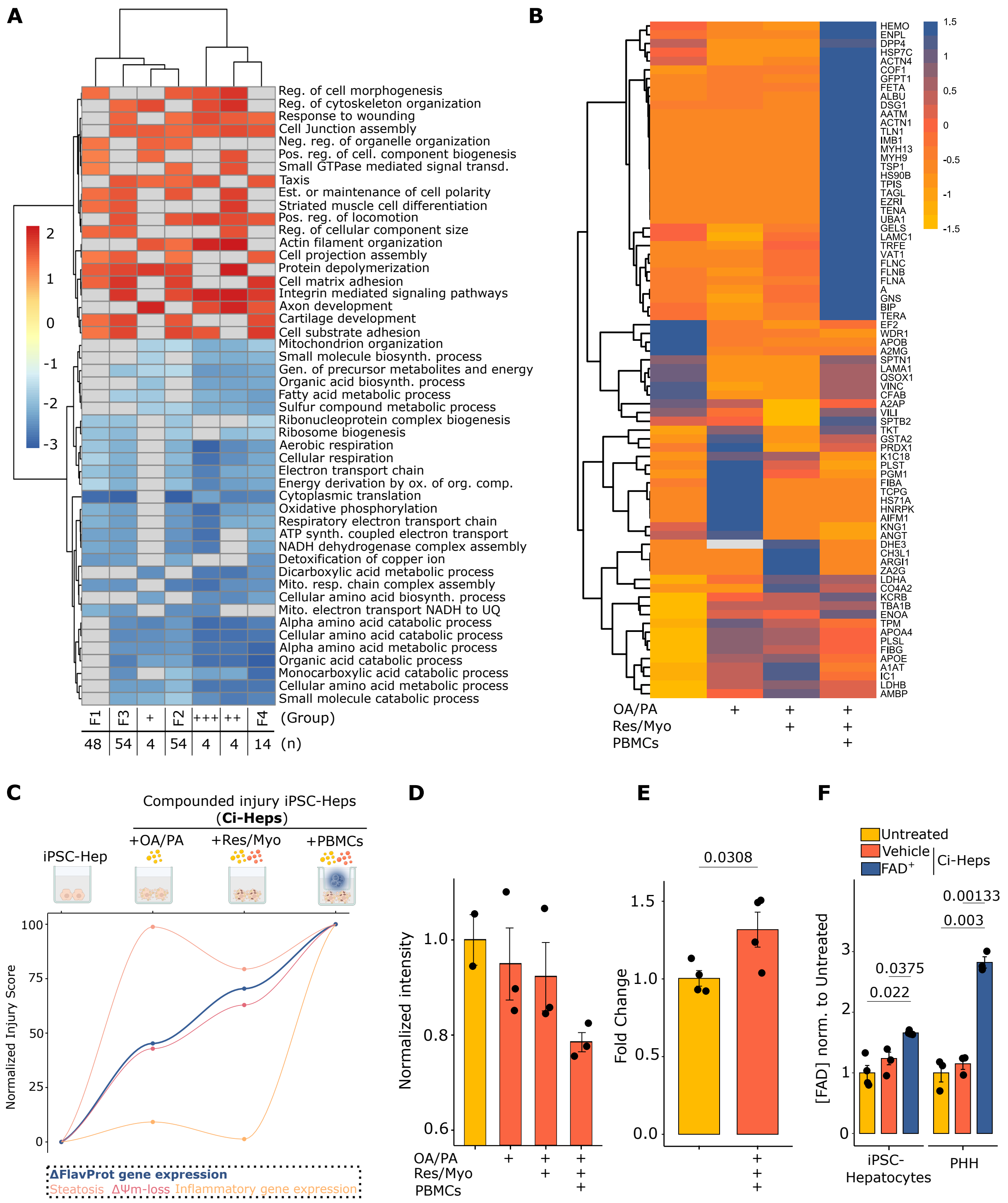

**Supp. Fig. 4: The flavoproteome (FlavProt) and its associated processes are dysregulated in MASLD models. A)** Clinical benchmarking of the *in vitro* MASLD models to clinical MASLD stages from Goavaere et al. stratified by fibrosis using Gene Set Enrichment Analysis (GSEA)-based clustering. GSEA was performed for all *in vitro* MASLD conditions compared to the untreated control. For the clinical dataset, all stages were compared to stage F0. **B)** Heatmap of the mean intensity-based absolute quantification (IBAQ) values of the identified human proteins. Proteins were identified in the supernatants of untreated induced pluripotent stem cell derived (iPSC-) hepatocytes or exposed to different stimuli (oleic and palmitic acid (OA/PA), resistin and myostatin (Res/Myo), co-cultured peripheral blood mononuclear cells (PBMCs)) using liquid chromatography-tandem mass spectrometry (LC-MS/MS). Heatmap shows the mean IBAQ values of n = 2 independent samples (except for untreated cells (n=1)) (see **Supp. Table 4**). **C)** MASLD injury index in iPSC-Ci-Heps exposed to different MASLD models. All parameters were standardized, ranging from untreated (0%) to the most complex disease condition including all disease triggers (100%). Inflammatory (Infla.) gene expression includes the genes *TNFA*, *CXCL8*, *CXCL1* and *CCL2*. FlavProt gene expression includes the changes for all genes from the human FlavProt. $\Delta\Psi m$ = mitochondrial membrane potential **D)** $\Delta\Psi m$ changes across iPSC-Ci-Hep conditions in another donor line **E)** Gene expression levels assessed in iPSC-Ci-Heps by qPCR for the *RFK* gene. P-value is shown after applying a one-sided t-test, as directionality was hypothesized based on the results from two other iPSC-Ci-Hep lines **F)** FAD levels in control and Ci-Heps for iPSC-derived and primary human hepatocytes. P-values are shown after applying multiple t-test, corrected using the Benjamini-Hochberg method.

Supplementary Figure 5:

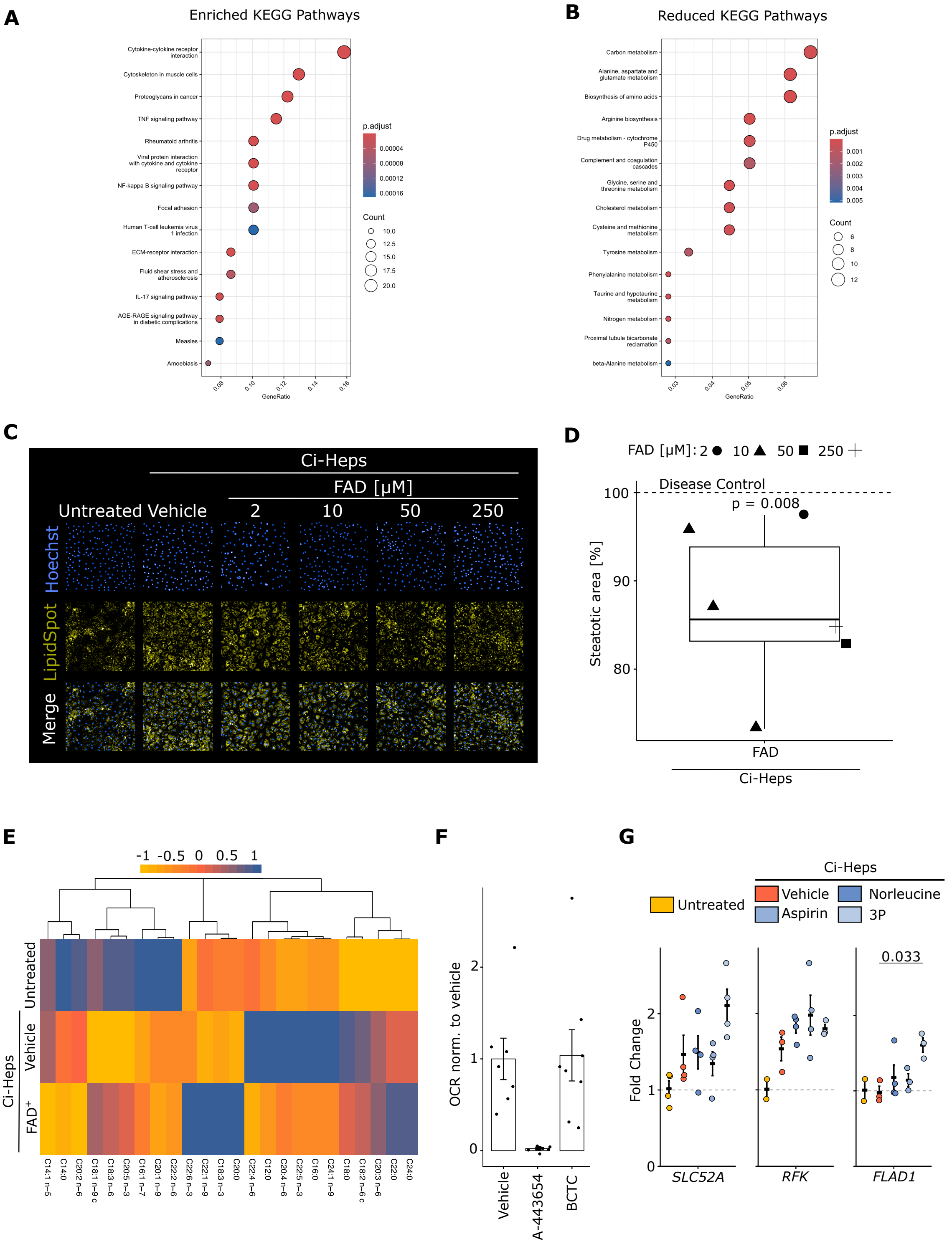

**Supp. Fig. 5: The flavoproteome (FlavProt) is a tractable target to intercept MASLD in hepatocytes.** KEGG Pathway enrichment analysis for the up- (**A**) and down-regulated (**B**) genes in induced pluripotent stem cell-derived compounded injury hepatocytes (iPSC-Ci-Heps) undergoing oleic and palmitic acid (OA/PA), resistin and myostatin (Res/Myo) and co-cultured peripheral blood mononuclear cell (PBMCs) exposure relative to the untreated control. **C)** Representative images of fluorescent staining of lipid droplets and nuclei in untreated iPSC-Heps or iPSC-Ci-Heps exposed to OA/PA + Res/Myo treated with different flavin adenine dinucleotide (FAD) concentrations. **D)** Quantification of the steatotic area normalized to the nucleus count in iPSC-Ci-Heps from C). Significance was used using a one-sided one sample t-test using the 100% vehicle control marked by the dotted line as a reference. Each dot represents an independent sample. **E)** Lipidomic heatmap analysis of iPSC-Heps or iPSC-Ci-Heps exposed to Res/Myo and trans-well PBMC co-culture with or without FAD supplementation (50 µM). Fatty acid species abundance was normalized by protein concentration and to total amount of measured fatty acid content. Heatmap shows the mean of n=4-6 independent samples. **F)** Spare respiratory capacity assessed via Seahorse in primary murine hepatocytes exposed to OA/PA and Res/Myo with or without candidate molecules. OCR = Oxygen consumption rate **G)** Gene expression changes assessed by qPCR for the vitamin B2 homeostasis genes in iPSC-Heps or iPSC-Ci-Heps exposed to OA/PA, Res/Myo and PBMCs with or without predicted FlavProt-active compounds. Fold change was assessed relative to the untreated control. Each data point represents an individual sample. Significance was assessed using multiple t-test corrected by the Benjamini-Hochberg method. Non-significant changes are not shown. 3P = 3-Pyridin-4-yl-1H-indazol-5-ylamine dihydrochloride.

Supplementary Figure 6:

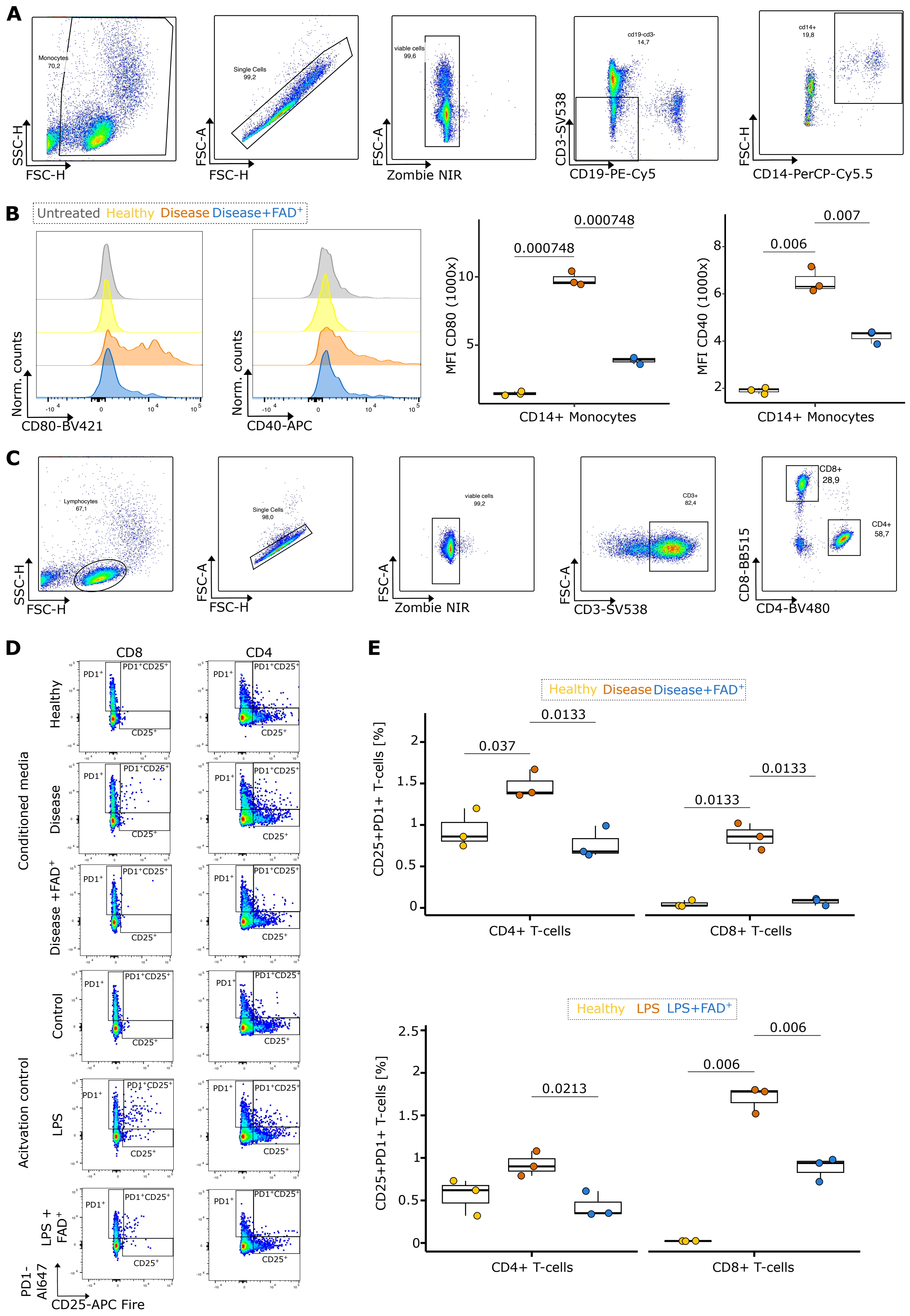

**Supp. Fig. 6: Characterizing anti-inflammatory effects by FAD supplementation (FAD$^+$) in primary human immune cells. A)** Full spectrum flow cytometry gating strategy for CD14$^+$ monocyte characterization in peripheral blood mononuclear cells (PBMCs) isolated from a healthy individual **B)** Representative histogram plots for the activation marker CD40 and CD80 in CD14$^+$ monocytes after exposure to supernatant from induced pluripotent stem cell (iPSC-) derived hepatocytes (Healthy) or iPSC-hepatocytes exposed to compounded injury (Disease) with or without FAD$^+$. **C)** Quantification of the parameters shown in B). Multiple t-test to assess significance, corrected according to the Benjamini-Hochberg method. **D)** Representative dot plots of T-cell activation marker in CD4$^+$ and CD8$^+$ T-cells stimulated with supernatant from iPSC-hepatocytes (Healthy) or iPSC-hepatocytes exposed to compounded injury (Disease) or LPS exposure with or without FAD$^+$. **E)** Quantification of T-cells double positive for the activation marker PD1 and CD25 after exposure to supernatants or LPS with or without FAD$^+$. Multiple t-tests to assess significance, corrected according to the Benjamini-Hochberg method. Non-significant p-values are not shown.

Supplementary Figure 7:

A

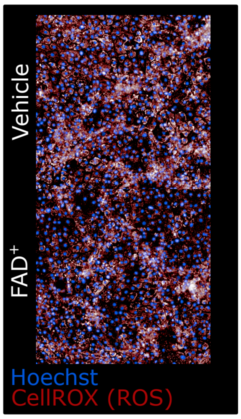


**Supp. Fig. 7: Assessement of reactive oxygen species (ROS) in hepatocytes undergoing metabolic stress. A)** Representative images of ROS in induced pluripotent stem cell derived hepatocytes undergoing compounded injury (oleic and palmitic acid and peripheral blood mononuclear cell co-culture) with or without supplementation of FAD ($FAD^{+}$).

Supplementary Materials and Methods

Drug prioritization strategies

Our computational drug prioritization pipeline has three individual strategies (**Table 1**): (1) Generation of a metabolic dysfunction-associated steatotic liver disease (MASLD-) specific biomedical knowledge graph (MASLD-KG), (2) Weighted gene co-expression network analysis of fibrosis stages in MASLD patients and (3) Analysis of single cell and single nucleus MASLD patient datasets to identify genes enriched in early staged diseased hepatocytes. We implemented approaches (2 and 3) using a published clinical MASLD dataset (Govaere et al., 2020[1]). For the following sections and the results, we utilize the new nomenclature of MASLD. Thus, early MASLD, refers to NAFL, MASLD to NAFLD and MASH to NASH in the respective data sets that use the old nomenclature as identifier.

**Table 1: Computational drug repurposing pipelines**

| **Approach** | **Sequence of analysis** |
|---|---|
| Knowledge Graph approach | |
| WCGNA approach | 1. MASLD Bulk RNA (fibrosis stages)<br>2. Protein-protein interaction networks<br>3. Drug-disease proximity analysis |
| Single cell approach | 1. MASLD Bulk RNA sequencing (early-stage vs control)<br>2. Single cell and single nucleus RNA-sequencing datasets<br>3. Protein-protein interaction networks<br>Drug-disease proximity analysis |

Generation of a MASLD biomedical knowledge graph

All analyses were performed on PrimeKG, a multimodal biomedical knowledge graph integrating 20 curated biomedical databases; the full graph comprised 129,375 nodes and 10,791,019 edges and was loaded as an undirected graph in python-igraph (v1.0.0).

Edge Filtering

CONTRAINDICATION and SIDE_EFFECT relationships (95,459 edges) were removed from the graph before any computation because both edge types connect drugs to diseases densely while carrying the opposite sign of the relationship being modelled. We augmented the PrimeKG disease-gene edges with two rule-based inferences, anatomy-mediated and phenotype-mediated. After rule evaluation (see below) removed the anatomy edges, and the graph was simplified to remove parallel edges and self loops, the reference graph contained 129,375 nodes and 3,975,530 edges.

Evaluation of the disease-gene inference rules

Both inference rules were validated on the ten diseases with at least 50 phenotype-rule genes (150–219 per disease; “MASH” itself had too few and was excluded, so rule decisions were generalised from these diseases). For each, we retrieved overall gene–target association scores from the Open Targets Platform (top 1,500 targets) and computed the ratio of the mean score of each rule's genes to that of degree-matched random gene sets (30 permutations), benchmarked against curated ASSOCIATED_WITH edges. The phenotype rule scored 2.5-fold above random (range 1.1–5.4; curated reference 5.0-fold) and was retained; the anatomy rule was indistinguishable from random (0.85-fold, range 0.32–1.02) despite contributing 4,921,285 edges (46% of the graph) and was removed.

Disease subnetwork extraction (MASLD-KG)

Starting from the “MASH” disease node, we extracted the union of all shortest paths of length one to three edges terminating at any drug node, giving a therapeutically oriented subnetwork rather than an undifferentiated three-hop neighbourhood. A forward breadth-first search builds the shortest-path directed acyclic graph, and a node is retained if it survives a backward closure from the drug targets. This MASH-seeded (seed-only) network contains 7,891 nodes and 506,612 edges. To incorporate disease context, we added differentially expressed genes from the GSE135251 liver-transcriptome cohort (|log2 fold change| > 2, adjusted $p < 0.05$) as additional seed nodes and expanded them by the same procedure. Of the resolved genes, 198 mapped to graph nodes; the DEG set is separate from the pre-specified target gene list (known positives), so the evaluation is independent of graph construction. The resulting graph, the MASLD-KG, contains 16,760 nodes and 1,759,226 edges, and is a strict superset of the seed-only network (all 7,891 nodes retained, 8,869 added).

Definition of MASLD-relevant pathway terms

MASLD relevance was measured against PrimeKG's own functional annotation, which derives from Reactome and the Gene Ontology and is independent of the GSE135251 transcriptomic signature used to build the graph, so the enrichment test is not circular. Six term families were specified *a priori* from established MASLD pathophysiology (lipid and fatty acid metabolism; fibrosis and extracellular matrix; inflammation; insulin and glucose handling; bile acid metabolism; oxidative stress), each defined by a case-insensitive regular expression applied to the names of all Pathway and BiologicalProcess nodes (31,158 terms). For example, the lipid family matched the terms lipid, fatty acid, triglyceride, cholesterol, lipoprotein, sterol and lipogen. This matched 1,426 unique terms (1,439 family assignments, since 13 terms belong to more than one family: inflammation 579, lipid and fatty acid 570, fibrosis and ECM 114, oxidative stress 81, insulin and glucose 66, bile acid 29). A gene was defined as MASLD-annotated if connected to at least one matched term, giving 4,403 of 27,671 genes (15.9%) as the background frequency. For a given gene set, fold enrichment was the number of MASLD-annotated genes observed divided by the number expected under the graph-wide annotation frequency (15.9%; 4,403 of 27,671 genes), with significance assessed by the hypergeometric test. Because fold enrichment against a fixed background is not comparable across gene sets of different size, enrichment was additionally expressed relative to a null of 200 random gene sets matched to each set on both size and log-degree profile, and graphs were compared only within size tier. The full list of matched terms with family assignments is provided as a supplementary file (Supplementary_mash_term_families.csv).

Topologically matched control diseases

To test whether the enrichment is specific to MASLD rather than a property of any central disease node, ten control diseases were matched to MASH in two stages. A first pass scored all 17,080 disease nodes on three-hop neighbourhood scale and node-class composition, using principal-component whitening and Mahalanobis distance to rank candidates by similarity to “MASH”. For each candidate the drug-reachable subnetwork was then extracted by the identical rule used for “MASH”, and matching assessed on the order, size and node-class composition of the subnetwork actually analysed. Candidates whose subnetwork had node-set Jaccard similarity above 0.95 to an already-selected control were rejected as effectively duplicate graphs, and diseases sharing MASLD biology were excluded. The ten controls (934 to 1,305 genes, spanning the 1,063-gene “MASH” subnetwork) were Kennedy disease, type 1 diabetes mellitus, ovarian endometriosis, sarcomatoid carcinoma, coronary artery disease, glomerulonephritis, congenital diaphragmatic hernia 1, salivary gland basal cell adenocarcinoma, undifferentiated carcinoma and androgenetic alopecia.

Size- and degree-matched enrichment

Fold enrichment against the whole-graph background is not comparable across subgraphs of different size, because larger neighbourhoods reach into less disease-specific biology. Each gene set was therefore additionally scored against a null of 200 random gene sets matched to it on both set size and log-degree profile, drawn from decile bins of the filtered graph, and enrichment is reported as the ratio of observed fold enrichment to the null mean. Graphs were compared only within size tier: seed graphs (934–1,305 genes) against one another, and augmented graphs (~5,500 genes) against one another. Augmentation controls were constructed identically to the MASLD signature, matched on gene count and log-degree profile, for both degree-matched random gene sets (n = 10) and signatures from ten unrelated diseases drawn from Open Targets (n = 10).

Seed-relative drug proximity

Global centrality measures importance of a node within a graph, not relevance to a seed disease, so drugs and genes were scored by random walk with restart (restart probability 0.15) seeded on the disease node. Because raw walk scores favour network hubs and are not comparable between graphs of different size, each score was converted to a z-score against a degree-matched null computed inside its own graph: for each permutation a random seed set of the same size was drawn from nodes of the same type and log-degree bin, and the null was built from 100 permutations network-wide, increased to 500 for the prioritised candidate drugs (random seed 42). “MASH” is zero-shot in this graph (no INDICATION edges, only 13 contraindications, which are excluded), so the method was validated separately on diseases with real indication data by held-out recovery.

Centrality and specificity

PageRank and exact betweenness were computed for every network in one process with one implementation (python-igraph, single-threaded), so values and runtimes are comparable. Exact betweenness (Brandes' algorithm) required 3,781 s (63.0 min) on the full PrimeKG graph against 113 s (1.9 min) on the MASLD-KG, a 33.4-fold reduction; total runtime fell 18.8-fold, from 3,782 s (63.0 min) to 201 s (3.4 min). Raw centrality is not comparable across networks of differing size, so each node's value was converted to a percentile rank within its own node class and network and compared across networks only where reference populations were of comparable size.

Software and reproducibility

Analyses used Python 3.14.2 with python-igraph 1.0.0, NumPy 2.3.5, pandas 2.3.3, SciPy 1.16.3 and statsmodels 0.14.6. All random procedures were seeded for reproducibility: the degree-matched proximity null used seed 42, and the random-gene augmentation control used seed 20260806.

Re-analysis of a bulk RNA-sequencing MASLD dataset:

Publicly available bulk RNA sequencing data from frozen liver biopsies were downloaded from the NCBI GEO repository (Home - GEO - NCBI (nih.gov), accession: GSE135251[1]). This dataset comprises 10 healthy controls, 51 early MASLD samples, and 155 MASH patients with varying stages of fibrosis. To analyze this dataset, we utilised the Nextflow workflow system. We first used nf-core/fetchngs v 1.10.0 to fetch the metadata and raw FastQ files from the Gene Expression Omnibus. We then used nf-core/rnaseq (version 3.12.0) to analyse the RNA sequencing data and produce a gene expression matrix. Here we utilized STAR to map the raw FastQ reads to the reference genome (GRCh38.p14.) and project the alignments onto the transcriptome followed by Salmon for quantification. Differential expression analysis (DESeq2) was performed comparing early MASLD and healthy samples with an absolute log fold change greater than 1 and a multiple testing corrected p-value lower than 0.05 was deemed significant. Visualizations included volcano plots, MA plots, and heatmaps of the top DEGs.

Weighted gene co-expression network analysis segmented by MASH fibrosis stage:

Gene co-expression network analysis was performed using the WGCNA R package[2,3] (version 1.73). Since the aim of this work was to construct disease specific co-expression network, the non-MASLD control samples were not included in this analysis.

First, the co-expression similarity matrix, calculated using a Pearson correlation coefficient for all possible pairs of genes, was transformed into an adjacency matrix using a soft-thresholding power β such that the resulting network structure satisfied the scale-free topology criterion. For this study, $\beta = 28$ was chosen. The adjacency matrix was then converted to a topological overlap matrix (TOM). Co-expression module identification was accomplished with the function cutreeDynamic from the dynamicTreeCut R package (version 1.63) using TOM dissimilarity (1-TOM) as the distance measure and a minimum module size cutoff of 30. Genes that were not assigned to any co-expression module were located in the 'grey' module. The resulting co-expression modules contained genes that were highly interconnected (co-expressed with each other).

To evaluate the correlation of co-expression module to fibrosis stage, the 216 MASLD samples were divided into the following subgroups using the clinical information provided by Govaere et al.[4]: early MASLD (51 samples), MASH F0-F1 (34 samples), MASH F2 (53 samples), MASH F3 (54 samples) and MASH F4 (14 samples), where 'F' indicates the degree of fibrosis in each group, while early MASLD patients displayed no fibrosis. We then converted the fibrosis stage metadata into a continuous numerical vector. Specifically, samples with no fibrosis were assigned a value of 0, samples with stage 1 fibrosis were assigned a value of 1, and so forth up to stage 4. The moduleEigengenes function (WGCNA R package) was used to calculate module eigengenes.

We then calculated the Pearson correlation coefficients between module eigengene expression and fibrosis stage, as well as its statistical significance, using functions from the WGCNA package. This approach allowed us to quantify the strength and direction of the association between gene module expression and fibrosis severity. Modules were considered significantly associated to fibrosis if p value $< 0.05$.

Pathway enrichment analysis of WGCNA modules

We performed overrepresentation analysis (ORA) to identify the main biological pathways representative of each module. First, the ensembl IDs for the genes in each co-expression module were converted to entrez IDs using the biomaRt R package (version 2.28.0). Enriched Gene Ontology terms, KEGG and Reactome pathways were then identified using the gProfiler2 R package (version 0.2.2). Enriched pathways were defined as those with FDR-adjusted p value < 0.05.

WGCNA-guided disease network construction

After having identified disease-relevant co-expression modules via WGCNA, we proceeded to expand such networks using the Dijkstra algorithm to identify all shortest paths between any two genes in a selected co-expression module. Each selected co-expression module was analyzed separately. This approach ensured that the resulting drug predictions could be directly linked to a biological process affected in MASLD.

First, the module genes were mapped to the human interactome. The human interactome was assembled from the following databases: STRING, IntAct, BioGRID, HI-Union, Interactome3D, HINT, APID, BioPlex3.0, SignaLink3.0. Were necessary, the databases were filtered to: include only interactions between human proteins, include only interactions validated experimentally and remove self-loops (protein interaction with itself. For data obtained from the string database, we also removed all interactions with confidence score < 0.4. This resulted in a network of 18748 proteins and 773143 edges. All shortest paths between all pairs of module genes in the human interactome were found using the all_shortest_paths function from the NetworkX python package (version 3.0).

Single-cell and single-nucleus RNA-seq MASLD datasets

To refine and validate the hepatocyte specificity of the bulk RNA-seq signature, four single-cell or single-nucleus RNA-seq datasets were analyzed: GSE210501, GSE166504[5], GSE212837[6], and GSE189600[7]. These datasets include mouse models of MASLD as well as human liver samples at various stages of disease. Each dataset was processed using the Seurat package (v4.3.0) following standard quality control procedures, including filtering of cells with high mitochondrial content or low feature counts. Normalization, identification of highly variable features, PCA, UMAP dimensionality reduction, and clustering were performed. Hepatocytes were identified based on canonical markers such as human and mouse genes for Alb, Cyp2e1, and Ttr. For each dataset, the refined MASLD gene signature was tested for differential expression within hepatocyte populations comparing disease versus control conditions. AUC analyses and UMAP-based visualizations were used to assess the discriminatory power of the signature, and expression scores were computed based on average expression of the upregulated targets.

Protein-protein interaction (PPI) network construction in MASLD samples:

To identify an early MASLD signature, we restricted the analysis of the GSE135251[1] dataset to early MASLD patients and healthy control samples. MASH patients were not considered as we wanted to identify the early changes, prior to the onset of fibrosis and inflammation.

To prioritize functionally central regulators within the MASLD transcriptional landscape, we constructed a protein–protein interaction (PPI) network using STRING v11.0 (combined score > 400). The input consisted of upregulated hepatocyte-specific genes derived from the bulk analysis. Shortest paths were computed between all gene pairs, yielding a high-confidence undirected network. Ensembl protein identifiers were converted to HGNC gene symbols using a curated alias dictionary. Subcellular localization data from the COMPARTMENTS database were used to annotate membrane and nuclear-associated genes.

Topological analysis of the PPI and WCGNA disease networks

Eigenvector centrality, degree, betweenness centrality and random walk with restart (RWR) were utilized to identify key proteins in the MASLD PPI network and in each WCGNA disease module. To identify significant key proteins, permutation tests were performed 1000 times for each of the network centrality algorithms. Each test generated a random network with a preserved degree distribution of the original disease network. These cumulative results were used to calculate the empirical p-value of the network algorithm. Proteins with empirical p value < 0.001 (from WGCNA module analysis) and empirical p value < 0.01 (from the single cell approach analysis) for either centrality measure were considered key nodes. The R and Python igraph packages were used for topological analysis respectively.

Drug-disease proximity analysis

FDA approved compounds and their interactions were sourced from ChEMBL[8], DrugBank[9], STITCH[10] and the Barabási group[11]. The network-based distance between drug targets and key disease proteins in the human interactome was calculated as described by Guney et al[12] for the WGCNA and Menche et al[13] for the single cell-based PPI analysis. Given K, a set of disease proteins, T a set of drug targets, and d(K, T), the shortest path length between nodes k ∈ K and t ∈ T in the human interactome, we define:

$$d_c(K,T) = \frac{1}{||T||}\sum_{t\in T} min_{k\in K} d(K,T) \quad (1)$$

The closest distance measure d(K, T) (average shortest path length between any drug target and its closest disease target, eq. 1) was used here because it showed the best performance in drug-disease pair prediction in the study by Guney et al[12].

To assess the statistical significance of the distance between a key protein and a drug dc(K, T), we performed 1000 permutation tests, each time with two randomly selected gene sets. Selection of random gene sets was performed so that the degree distribution of the random sets was similar to the real sets. However, due to the scale-free nature of biological networks, there are only a few high-degree nodes in the human interactome. To avoid repetitive selection of the same, high degree nodes during random selection, we used a binning approach in which nodes within a certain degree interval were grouped together such that there were at least 100 nodes in each bin. When selecting the random gene sets, we sampled nodes from the same bins as the real drug/disease targets. The mean and standard deviation of the reference distribution was then used to calculate the Z-score of the observed distance. Drugs with Z-score <= -1.96 (for WGCNA-driven drug discovery) and Z-score <= -2 (for PPI-driven drug discovery) were considered significantly proximal.

Network visualization and annotation

Final network visualisations were generated using Cytoscape (v3.9.1), iGraph and NetworkX (v2.6.3) with Force Directed, Fruchterman–Reingold and circular layouts. In the WGCNA network modules **(Supp. Fig. 2)**, nodes were colored by differential expression and betweenness centrality respectively. Differential expression was conducted using edgeR (v4.4.2) comparing all MASLD to control cases. Betweenness centrality computation was performed in Cytoscape. Figures were rendered at high resolution using CairoSVG and Matplotlib.

Human stem cell lines

We used 4 human induced pluripotent stem cell (iPSC-) donor lines in total. These included BIHi001-B major homozygous for *PNPLA3* rs738409 variant (*PNPLA3*- I148I) and BIHi005-A minor homozygous for the *PNPLA3* rs738409 variant (*PNPLA3*-I148M) provided by the BIH Core Unit Pluripotent Stem Cells and Organoids (CUSCO). Both cell lines are registered in the Human Pluripotent Stem Cell Registry (RRID:CVCL_YN70 and RRID:CVCL_IT56). Further, we used the BIHi314-A cell line derived from a MASLD patient with zone 3 MASH generated by the CUSCO and the FSPS13B hiPSC line for hepatocytes generated by Forward programming (Fop-Hep) as described in Tomaz et al[14]. We cultured all cells on vitronectin-coated plates (Thermo). BIHi001-B, -005-A and -314-A cells were cultured in mTeSR medium (Stem Cell Tech), FSPS13B cells were cultured in E8 (produced by the CUSCO). All experiments were performed at 37°C in a 5% $O_2$ and 5% $CO_2$ hypoxic atmosphere.

Differentiation into stem cell derived Hepatocytes

For the non-FoP-Hep differentiations we adapted a 28-day protocol[15], optimized by Seifert & Ludwik et al[16], and modified by us for the maturation of Hepatocytes, as following:

Generation and thawing of ROCKi-Adapted iPSCs:

iPSCs were expanded in mTeSR medium (StemCell Technologies) containing 10 µM ROCKi (Tocris), detached with 0.5 mM EDTA in PBS for 5 min at room temperature, resuspended in Bambanker freezing medium (NIPPON), frozen at -80°C using a Mr. Frosty container, and transferred to vapor-phase liquid nitrogen (-135°C). For thawing, vials were warmed in a 37°C water bath until a small ice crystal remained, diluted dropwise with 10× volume of mTeSR + ROCKi, centrifuged (3 min, 300 g), and resuspended in fresh ROCKi-containing mTeSR.

Hepatic Differentiation Protocol:

Upon reaching ~80% confluency, iPSCs were single-cell seeded onto Geltrex-coated 6-, 12-, or 24-well plates (Day 0) and differentiated as follows:

Days 0–5: DE-induction medium (RPMI1640, 2% B27, 1% PenStrep, 100 ng/ml Activin A, 50 ng/ml Wnt3, 0.5 mM sodium butyrate), followed by 1 day without sodium butyrate.
Days 7–12: KO-medium (KO-DMEM, 20% KO Serum, 1% PenStrep, 1% Glutamax, 1% NEAA, 1% DMSO, 0.1 mM β-mercaptoethanol) for hepatic commitment.
Days 13–19: HBM medium (Lonza) supplemented with 20 ng/ml HGF, 20 ng/ml OSM, and 1% PenStrep.
Days 20–28: Maturation in HCM Bulletkit medium (Lonza, excluding EGF), with supplementation of 5 nM T3 (Tocris), 100 nM sodium selenite (Sigma), and 100 µM CDCA (Sigma)[17,18].

Forward programming of FoP-Heps was performed based on Tomaz et al[13]. In brief, cells contain a doxycyclin-inducible expression system (Opti-OX) of *HNF1A, HNF6, FOXA3* and *RORCc*. The iPSCs were harvested as single cell suspension with accutase and 17,000 cells / cm$^2$ were seeded in 12 well format in E8 (produced by BIH-CUSCO), 1% PenStrep (Thermo), 10 µM Y-27632 2HCl / Rock inhibitor (Selleckchem) (day –1).
Day 0: E8 (produced by CUSCO), 1% PenStrep (Thermo).
Day 1: E6 (Gibco), 1% PenStrep, (Sigma Aldrich),1 mg/ml doxycycline (MP Biochemicals).
Day 2, 3, 4, 5, 7, 9 HepatoZYME-SFM (Thermo), 2% chemical defined lipid concentrate (Thermo), 2% MEM Non-Essential Amino Acids Solution (Thermo), 1% PenStrep (Thermo), 14 µg/ml insulin (Sigma-Aldrich), 30 µg/ml Human Apo-Transferrin, CF (R&D), 50 ng/ml HGF (Preprotec), 20 ng/ml OSM (R&D), 1 µg/ml doxycycline (MP Biochemicals.)
Day 10,12,14, 16, 18, 19 HepatoZYME-SFM (Thermo), 2% chemical defined lipid concentrate (Thermo), 2% MEM Non-Essential Amino Acids Solution (Thermo), 1% PenStrep (Thermo), 2 µg/ml insulin (Sigma-Aldrich), 30 µg/ml Human Apo-Transferrin, CF (R&D), 50 ng/ml HGF (Preprotec), 20 ng/ml OSM (R&D).

<u>Primary hepatocytes</u>
The primary human hepatocytes (PHHs) were isolated from individuals undergoing liver surgery due to colorectal liver metastasis. Cell isolation was done under informed consent and the ethics statement EA1/257/14 according to the protocol described in Horner et al[19]. PHHs were frozen in FCS with 10% DMSO using a Mr. Frosty container and stored in vapor phase liquid nitrogen below -135°C until further use. For the experiments using cells from the 63-year-old individual, we utilized a recently described protocol[20]. Briefly, cells were plated on collagen-coated plates (collagen type 1, rat tail, Gibco), a Matrigel overlay was applied, and cells were maintained in complete HCM medium (Lonza). For the cells from the remaining donors, we did not include a matrigel overlay and kept them in WilliamsE medium including 10% FCS, 1% Pen-Strep, 1% Sodium Pyruvate, 1% Glutamax, 1.5% Hepes buffer, 0.12 I.E./ml insuline and 0.8 µg/ml dexamethason.
Other primary hepatocytes were isolated from C57BL/6J mice (12–20 weeks old) using a well-established two-step perfusion technique. Initially, the liver was perfused with a calcium-free buffer to remove blood and non-parenchymal cells, followed by perfusion with collagenase (LiberaseTM, Roche, Switzerland) to dissociate the tissue enzymatically. The resulting cell suspension was purified using a Percoll gradient, yielding hepatocytes with >90% viability as previously described[21]. Purified cells were seeded onto collagen type I-coated culture plates (5 µg/cm$^2$) at densities suitable for experimental needs. Cells were maintained in William's E medium supplemented with 2 mM L-glutamine, 5% PenStrep, and hepatocyte growth supplements (CM4000; Thermo Fisher Scientific). This complete medium supported optimal growth and maintenance of hepatocytes *in vitro*.
All animal experiments complied with ethical regulations and were approved by the Committee on Research Animal Care of the Charité Universitätsmedizin Berlin, Berlin, Germany (License number: G0174/20).

<u>Primary human peripheral blood mononuclear cells (PBMCs)</u>
Blood isolation was done in accordance with ethical standards of the Charité Universitätsmedizin Berlin under the ethical approval EA2_065_21. We obtained healthy human PBMCs using Ficoll-Paque (Sigma). To minimize PBMC donor variability, we derived them from a standard "master" donor who a physician clinically monitored to ensure the absence of clinical illness or medication intake. Specifically, we mixed whole blood with PBS (w/o $Ca^{2+}$ $Mg^{2+}$) in a 1:1 ratio and overlaid this dilution on the Ficoll in a 3 to 2 ratio, respectively. We then centrifuged the mixture for 30 minutes at 300 g at room temperature, using the lowest levels of brake and acceleration. The leukocyte layer was transferred to a new tube and washed once with PBS. After lysing red blood cells by incubating the

cells for 10 min in ACK lysis buffer, we washed twice, counted and froze cells down in a 10% DMSO solution in FCS using Mr. Frosty. The cells were stored in vapor phase liquid nitrogen below -135°C until further use.

Compounded metabolic injury model

We tested different combinations of compounded metabolic injury hits in hepatocytes (Ci-Heps). Generally, we exposed iPSC-Ci-Heps for 3 days to either free fatty acids (FFAs) alone (1 Hit), combined with the inter-organ signaling molecules resistin (Res) and myostatin (Myo) (2 Hit) or in addition with co-cultured PBMCs (3 Hits). For rescue experiments, we also included combinations as indicated.

For FFA injury, we exposed cells to oleic and palmitic acid (OA and PA) in a 5 to 2 ratio on day 28 for 3 days. OA and PA (both Merck) were coupled to the HCM bullet kit-derived BSA by incubation at 37°C for at least 2 h. We used 125 µM as total fatty acid concentration.

To account for inter-organ signaling in our model, we exposed Ci-Heps to recombinant resistin and myostatin (100 ng/ml each, Peprotech) for three days.

For modeling inflammation, we included PBMCs in an indirect trans-well co-culture using Millicell trans-well inserts (Millipore) with a 0.4 µm pore size, 1 day after the addition of the other injuries (Day 29). We thawed the PBMCs 18 h before co-culture and cultured them in RPMI medium containing 10% heat-inactivated fetal calf serum (FCS) and Pen-strep. Within this 18 h pre-incubation, PBMCs incubated with 5 µg/ml lipopolysaccharide (LPS, eBioscience or Stem Cell Technologies). Before adding them to the inner compartments of the trans-wells, we washed off the RPMI medium and re-suspended them in the respective HCM culture medium. PBMC viability after coculture was consistently over 90%, as assessed via Trypan Blue staining. To derive an overall injury metric (**Fig. 3A**), we normalized each disease marker for all conditions to untreated (0%) and Ci-Heps exposed to 3 Hits (100%).

For the FoP-Heps we started disease modeling on day 13 or day 16, due to their faster differentiation. For the primary human hepatocytes, we applied recombinant TNFa and IL1B, 10 pg/ml each, instead of PBMC co-culture.

Live dyes

We incubated the cells for 30 min with Hoechst, LipidSpot 488/610 and MitoTracker Red CMXRos staining to assess steatosis and mitochondrial membrane potential. For measuring reactive oxygen species (ROS), we used CellROX Deep Red (Thermo). We used all dyes according to manufacturer´s instructions. Specifically, Mitotracker was used with a 200 nM end concentration. Eventually, we either performed live-cell imaging or fixed cells in the culture plate using 4% paraformaldehyde, followed by subsequent PBS washes before imaging.

Fluorescence imaging and image analysis

We imaged cells in their respective well plates using the Phenix High-Content Imaging system (Revity) or the Olympus CKX53. We captured at least 10 individual images per well on the Phenix, or 3 individual images per well on the Olympus, which we analyzed using the Signals Image Artist software (Version 1.3.29) or Fiji, respectively. For measuring steatosis, mitochondrial membrane potential, or ROS we manually set upper and lower thresholds for the respective staining in each experiment to eliminate background and artefacts. For Mitotracker and ROS stainings, we then measured intensity for the selected image region. To assess steatosis, we measured LipidSpot area and divided it by the respective Mitotracker area or number of nuclei for each image. Respective signals for steatosis or Mitotracker we then further normalized as indicated.

RNA Isolation

We isolated RNA using a hybrid protocol of trizol RNA isolation combined with column-based purification (Qiagen or Thermo). We lysed cells in 1 ml of trizol (Thermo) to which we subsequently added 0.2 ml chloroform. All chloroform-trizol mixtures were then centrifuged for 10 min, using 12,000 g at 4°C. Then, we transferred the aqueous phase to a new tube and added 70% ethanol in a 1:1 ratio. This mixture we loaded on RNeasy Micro Kit columns (Qiagen). From this point, we performed every step according to the manufacturer´s recommendations (including the on-column DNAse treatment). We assessed the concentration of the isolated RNA using Nanodrop.

qPCR

For quantifying gene expression using qPCR, we reverse-transcribed RNA into cDNA using a cDNA Synthesis kit (High-Capacity cDNA Reverse Transcription Kit (Thermo, Cat. No. 4368813) according to manufacturer´s instructions. We then performed RT-qPCR using the PowerUp SYBR Green Master Mix (Applied Biosystems, Cat. No. A25776) on a QuantStudio 3 Real-Time PCR System (Thermo Fisher Scientific Inc.) according to the manufacturer's instructions. Primers were synthesized by Eurofins Genomics, with sequences listed in **Table 2**. Gene expression changes were calculated using the ddCT method.

**Table 2: Oligos used in this study**

| | |
|---|---|
| *RFK*_F | GTGGTGAGCATAGGATGGAACC |
| *RFK*_R | GGTCTCAGGTAGCCAACAATGG |
| *RPS18*_F | GCAGAATCCACGCCAGTACAAG |
| *RPS18*_R | GCTTGTTGTCCAGACCATTGGC |

Supernatant analysis

We collected supernatant from cells exposed to the indicated disease models 18-24h after the last medium change, albeit consistent timing for individual experiments. We stored the samples at -80°C until further use. The molecules TNFa, IL1B and aspartate aminotransferase (AST) were measured by Labor Berlin–Charité Vivantes GmbH, Berlin, Germany, using standard clinical biochemical procedures that can be detailed upon request.

Conditioned media analysis using Liquide chromatography – tandem mass spectrometry (LC-MS/MS)

For conditioned media analysis, we analyzed the conditioned supernatant of cells using mass spectrometry. For more sensitive detection of secreted proteins, we added HCM medium without additional growth factors or BSA for 24 h after exposure to the respective disease model. We centrifuged the samples and stored the supernatant at -80°C until the analysis.

For the analysis, 18 µl of each culture supernatant were separated by one-dimensional SDS-polyacrylamide gel electrophoresis according to Beier et al.[22]. Each sample lane was divided into three sub-samples (all protein bands above the bovine serum albumin (BSA) band, the BSA band and all protein bands below the BSA band). In-gel digestion of proteins, extraction of peptides, desalting and preparation of the samples for LC-MS/MS analysis were performed as described by Beier et al [22]. The LC-MS/MS analysis was performed using a nanoAQUITY UPLC system (Waters corporation, Milford, MA, USA) coupled to an LTQ Orbitrap Velos Pro mass spectrometer (Thermo Fisher Scientific Inc., Waltham, Massachusetts; USA). The LC gradient used is described in Lerch et al[23]. A m/z of 350-1750 was used for primary MS scans. Except for the exclusion time, the other MS parameters were taken from Beier et al[22]. An exclusion time of 12 s was chosen. MS/MS raw files were analysed using MaxQuant (Max Planck Institute of Biochemistry, Martinsried, Germany, www.maxquant.org, version 2.4.3.0 according to the parameters described previously (Beier et al.[22]) using N-terminal acetylation as an additional variable modification against a database of human proteins (taxonomy id 3A9606_29) and isoforms downloaded from Uniprot on 2023-06-27 and a database of Bos taurus proteins downloaded from Uniprot on 2023-12-20. Proteins are considered to be reliably identified if they are identified by at least two unique peptides, each of them detected by at least one MS/MS scan in two different samples/replicates. In addition, their total intensity must be greater than 100.000. Protein quantification was performed using MaxQuant (version 2.4.3.0) intensity-based absolute quantification (iBAQ) (Schwanhäusser et al.[24]). The means of two biological replicates (for stem cell derived hepatocytes without any trigger only one replicate was measured) were normalised and plotted as a heatmap using the R package tidyHeatmap2 after subtracting the means of the blank samples (culture medium). The data input is provided in **Suppl. Table 4.** The mass spectrometry proteomic data have been deposited in the ProteomeXchange Consortium via the PRIDE partner repository (Perez-Riverol et al[25]) with the dataset identifier PXD063495. Reviewers will have access to the MS data at project accession: PXD063495 using the token lSRc2SuWKYkF. Alternatively, reviewer can access the dataset by logging in to the PRIDE website using the following account details: Username: reviewer, Password: 7jwiwNNKtUYa

Bulk RNA Sequencing

mRNA sequencing for characterizing the MASLD *in vitro* model was performed at the BIH Genomics Core facility using Illumina TruSeq stranded mRNA library preparation and for sequencing using 50 mln of PE100nt reads per sample on a 25B flowcell of the Illumina NovaSeq X Plus. For exploring transcriptomic changes by FAD supplementation (FAD rescue) we used the services from Novogene. The sequencing of the FAD rescue samples was performed using the PE150 sequencing strategy on NovaSeq Xplus sequencer 25B sequencing flow cell (Illumina, San Diego) by Novogene Germany (Munich, Germany).

Features of the MASLD *in vitro* model dataset:
The MASLD *in vitro* model bulk RNA-seq dataset consisted of 32 samples from two donor lines *PNPLA3*-I148I and -I148M (16 for each donor line, four of each sample), see **Table 3**.

**Table 3: Sample conditions used for bulk RNA-sequencing and downstream analysis for characterizing the *in vitro* MASLD models.**

| Name | Condition |
|---|---|
| HCM | healthy control (untreated cells) |
| OA PA | cells that were solely treated with an excess of free fatty acids |
| OA PA ResMyo | cells that in addition to free fatty acid excess also were exposed to 2 MASLD associated signaling molecules from other tissues (adipose and muscle tissue) |
| OA PA ResMyo PBMCs | cells that experienced both of the above-mentioned conditions + a co-culture of stimulated primary immune cells (immune cells were not included in the RNA sequencing) |

Processing and analysis of MASLD *in vitro* model RNA-seq dataset:
We started by carrying out a whole genome decoy-aware indexing and transcript quantification using Salmon v1.10.1 [26]. The Salmon pipeline estimates transcript-level abundance from RNA-seq read data. We then used DESeq2 [27], to test for differentially expressed genes between the following conditions:

- **I.** OA PA vs HCM
- **II.** OA PA ResMyo vs HCM
- **III.** OA PA ResMyo PBMCs vs HCM

For derivation of the totatl flavoproteome (FlavProt) gene expression change (**Fig. 3A**), the counts for each FlavProt gene[28] in each sample were normalized to the total counts and to its untreated control. The mean of the overall change per condition, independent of direction, was normalized to untreated (0%) and Ci-Heps exposed to 3 Hits (100%).

Signaling pathway enrichment in MASLD *in vitro* model dataset:
For each set of differentially expressed genes, we tested for enriched KEGG pathways[29].

Clinical benchmarking using gene set enrichment analysis:
To functionally characterize the *in vitro* models, we used gene set enrichment analysis (GSEA). GSEA is a computational method used to identify whether a set of predefined genes shows statistically significant differences in expression between two biological states (e.g., disease vs. control). Unlike traditional approaches that focus on individual genes, GSEA evaluates the expression of entire gene sets (groups of functionally related genes) to see if they are more likely to be upregulated or downregulated in a particular condition. In the context of this study GSEA helps in functional characterization by providing insights into the biological processes that are associated with the observed changes in gene expression. DEGs from each of the MASLD *in vitro* model datasets were subjected to gene set enrichment analysis (GSEA) using the “GSEA” function in R. The “msigdb” package in R was used to extract predefined gene sets (GO:BP) from the Molecular Signatures Database. Gene Ontology (GO) biological processes were enriched with a significance threshold of $p < 0.05$. This analysis was also performed for the DEGs from the Govaere et al. dataset[4], stratified by the fibrosis stages described in their study. The analyzed gene sets are summarized in **Table 4**.

**Table 4: Datasets used for clinical benchmarking**

| MASLD *in vitro* model | Govaere et al Fibrosis Stage |
|---|---|
| OA PA vs HCM | Control |
| OA PA ResMyo vs HCM | F1 |
| OA PA ResMyo PBMCs vs HCM | F2 |
| | F3 |
| | F4 |

Features of the FAD rescue dataset:
The FAD rescue bulk rna-seq dataset initially consisted of 8 samples (4x samples per condition) derived from the *PNPLA3*-I148I line which were lentivirally transduced with EGFP 2 days prior to the disease modeling (**Table 5)**. Due to poor RNA quality of one sample from the OA/PA + Res/Myo + PBMCs + FAD condition which did not pass the Novogene sample QC, we had to remove it from the analysis.

**Table 5: Sample conditions used for bulk RNA-sequencing and downstream analysis of the FAD rescue.**

| Name | Condition |
|---|---|
| OA PA ResMyo PBMCs | Ci-Heps exposed to the OA/PA, Res/Myo and PBMC disease model which were lentivirally transduced with EGFP |
| OA PA ResMyo PBMCs + FAD | Ci-Heps exposed to the OA/PA, Res/Myo and PBMC disease model which were lentivirally transduced with EGFP and treated with 50 μM of FAD |

The processing and data preparation was carried out by Novogene´s bioinformatic services. The generation of DEGs which were also used as an input for KEGG pathway analysis we carried out using the Novogene magic web application for the following comparisons:

**I.** OA PA ResMyo PBMCs vs OA PA ResMyo PBMCs + FAD

To assess the directonality FlavProt gene expression under injury and under rescue (**Fig. 3B**) we calculated for both RNA seq experiments the relative expression changes of each gene between OA PA ResMyo PBMCs to Untreated for the available data from both cell lines, and OA PA ResMyo PBMCs to OA PA ResMyo PBMCs + FAD. Genes without detectable counts were removed.

Quantification of intracellular FAD
We quantified intracellular FAD levels using the fluorescence-based detection of the FAD assay kit (Abcam, ab204710) according to the manufacturer's instructions. Before harvesting cells for the assay, we washed 3x times with PBS to ensure no significant amounts of exogenous FAD remained. We performed the assay read-out using the Spectramax i3X Plate reader.

Identification of compounds with similar network topology to FAD
Given the impact of mitochondrial dysfunction in MASLD[30], we refined MASLD signatures to a subset of the MASLD DEG samples associated with mitochondrial structure or function. The list of mitochondrial-associated genes was sourced from the MitoCarta 3.0 database[31]. We then selected compounds with shared nodes to FAD. We further expanded the drug search to include 24,000 investigational and experimental compounds in phase 1-3 clinical trials **(Supp. Table 6)**. From these, we prioritized norleucine (Nor), aspirin (Asp), 3-Pyridin-4-yl-1H-indazol-5-ylamine dihydrochloride (3P), BCTC (Sigma) and A-443654 (Biomol) for rescue experiments.

Rescue experiments for additional compounds

For the rescue experiments, we co-exposed the cells to the indicated concentrations of FAD (Roth), Nor (1 mM; Thermo), Asp (5 µM; Sigma), 3P (1.5 µM; Sigma), BCTC (3 µM; Sigma) and A-443654 (5 µM; Biomol) plus the indicated injury condition using the duration and concentrations described for the Ci-Heps. Thus, the total exposure to each compound was 72 h. After the co-exposure, we harvested the respective material for the analysis as indicated.

Measurement of the oxygen consumption rate (OCR)

For testing the rescue effect by FAD or other compounds, we co-exposed primary mouse hepatocytes to the compound plus OA/PA and Res/Myo for three days. We measured OCR using the Seahorse Extracellular Flux (XF)-96 Bioanalyzer from Agilent Technologies, as described previously[32]. Briefly, the XF basal medium from Agilent Technologies was supplemented with 10 mM glucose, 1 mM pyruvate, and 1 mM glutamine. The electron transport chain complexes inhibitors (oligomycin (2 µM), FCCP (1 µM), rotenone (1 µM), and antimycin (1 µM)) were titrated and delivered through injection at designated ports A, B, and C of the Bioanalyzer. The resulting data were normalized to the protein content in each well using the Pierce™ BCA assay (Thermo Fisher Scientific).

Measurement of electron transport chain complex activities via respirometry-based assay

The electron transport chain complex I, II, and IV activities were measured in frozen hepatocytes using Seahorse XF-96 Extracellular Flux Bioanalyzer (Agilent Technologies, Santa Clara, CA, USA) as described[33]. Primary mouse hepatocytes were treated with the respective treatments for 72 hours. After treatment, the medium was removed, cells were washed with PBS three times. Then, the cells were harvested and frozen at -70°C. On the day of assay, cells were homogenized in MAS buffer (70 mM sucrose, 220 mM mannitol, 5 mM $KH_2PO_4$, 5 mM $MgCl_2$, 1 mM EGTA, 2 mM HEPES, pH 7.4) using 10 strokes in a glass Dounce homogenizer. The homogenate was centrifuged at 200 g for 10 minutes at 4°C, and the supernatant was collected. Protein concentration was measured using the Pierce™ BCA assay (Thermo Fisher Scientific).

For the Seahorse XF-96 assay, 15 µg of homogenate protein was loaded into each well in 20 µl of MAS buffer. The plate was centrifuged at 2,000 g for 5 minutes at 4°C, after which 130 µl of MAS buffer containing cytochrome c (10 µg/ml) was added to each well. Substrates were delivered through port A (NADH, 1 mM; or succinate + rotenone, 5 mM + 2 µM). Inhibitors rotenone and antimycin A (2 µM and 4 µM, respectively) were delivered through port B, TMPD + ascorbic acid (0.5 mM + 1 mM) through port C, and azide (50 mM) through port D. The data was normalized to the proteins content. OCR signal after azide injection was subtracted from NADH (complex I activity), succinate + rotenone (complex II activity), and TMPD+ascorbic acid (complex IV activity).

Lipidomics

We harvested Ci-Heps that we left untreated or exposed to Res/Myo and PBMCs with or without 50 µM of FAD. We omitted exogenous FFAs to focus on cell-intrinsic changes. After exposure, we harvested the cells by washing them twice with PBS and subsequently incubating them in 10X TryPLE. After the cells dissociated from the dish, we inactivated TryPLE using PBS with 20% KO-Serum and centrifuged them for 3 min using 200 g. The Pellets were then stored in -80°C for 24 h until the analysis. For the analysis cells were hydrolyzed with 100 µL 10 mol/L NaOH within 120 min at 80 °C. The samples were neutralized with 100 µL acetic acid and filled up to 5 mL with methanol containing internal standards (Cayman Chemical, Ann Arbor MI). HPLC measurement was performed using an Agilent 1290 HPLC system with binary pump, autosampler and column thermostat equipped with a Phenomenex Kinetex-C18 column 2.6 µm, 2.1 x 150 mm column (Phenomenex, Aschaffenburg, DE) using a solvent system of acetic acid (0.05%) and acetonitrile. All solvents and buffers in LC-MS-grade were purchased from VWR Germany. The solvent gradient started at 70 % acetonitrile and was increased to 98 % within 10 min and hold until 14 min with a flow rate of 0.4 mL/min and 5 µL injection volume. The HPLC was coupled with an Agilent 6470 triplequad mass spectrometer with electrospray ionisation source operated in negative selected ion mode.

FAD rescue in primary immune cells

We freshly isolated healthy PBMCs as described above, used for disease modeling, and cultured 50000 cells in 100 µl of RPMI with 10% heat-inactivated FCS and Pen-Strep in a 96-well plate. To these, we either added 50 µl of supernatant from healthy iPSC-hepatocytes or from Ci-Heps exposed to OA/PA, Res/Myo and PBMCs with or without freshly added FAD (50 µM). We included 5 µg/ml of

LPS (Stem Cell Technologies) as a positive control, with or without 50 µM FAD. After 18 h, we harvested the cells for full-spectrum flow cytometry, stained for the Panel in **Table 6**. Briefly, we transferred cells into FACS tubes and washed off the medium using FACS buffer (PBS with 2 mM EDTA). After centrifugation, we discarded the supernatant, and the viability dye was added to the residual volume. After 10 minutes, we added the antibody mix that we prepared in blocking buffer. After 20 min, we washed the samples using FACS buffer. Subsequently, we fixed the cells in 2% formalin for 10 min, which we washed off using FACS buffer. Then, samples were acquired on a Cytek® Aurora Cytometer with 3 lasers (Violet, Blue, Red) and 38 detectors with the Spectroflo® v3.0.3 software. Daily QC was performed according to the manufacturer's instructions. After unmixing, cells were analyzed with the gating indicated.

**Table 6: Flow cytometry-antibodies and dyes used in this study**

| Channel | Marker | Target | Clone | Manufacturer | Ordering No |
|---|---|---|---|---|---|
| BV421 | CD80 | Myeloid activation | 2D10 | BioLegend | 305222 |
| PacB | CD66b | Neutrophils | G10F5 | BioLegend | 305112 |
| BV480 | CD4 | CD4TC | RPA-T4 | BD | 746541 |
| V500 | CD16 | Neutrophils/monocytes | 3G8 | BD | 561394 |
| SV538 | CD3 | TC | UCHT1 | BioLegend | 300484 |
| BV570 | HLA-DR | MHC-II | L243 | BioLegend | 307638 |
| BV711 | CD40L | TC activation | 24-31 | BioLegend | 310838 |
| BV750 | CD56 | NK cells | 5.1H11 | BioLegend | 362556 |
| BB515 | CD8 | CD8TC | RPA-T8 | BD | 564526 |
| Alexa532 | CD45 | Pan-immune | HI30 | eBio | 58-0459-42 |
| PE-Cy5 | CD19 | B cells | HIB19 | BioLegend | 302210 |
| PerCP-Cy5.5 | CD14 | Monocytes | M5E2 | BioLegend | 301824 |
| PE-Fire810 | CD86 | Myeloid activation | IT2.2 | BioLegend | 305455 |
| APC | CD40 | Myeloid activation | 5C3 | BioLegend | 334310 |
| Al647 | PD1 | TC activation | NAT105 | BioLegend | 367420 |
| Al700 | CD11c | APC/DC | Bu15 | BioLegend | 337220 |
| Zombie NIR | Viability | | | BioLegend | 423106 |
| APC Fire 810 | CD25 | TC activation | M-A251 | BioLegend | 356150 |

Analysis of survival and liver-related events in adult MASLD cohort

We analyzed the SteatoSITEv2 cohort to hepatic events (decompensation) with FlavProt gene expression. Kaplan-Meier Plots were generated as described in Schwabe et al[34] and tested for significance using a log-rank test. We therefore performed lasso-penalised cox regression including gene expression of all FlavProt genes for all cases with successful RNA-seq where there was no decompensation coding event prior to the biopsy (n = 446). For the coefficients from the resulting gene signature, we derived a single metric value for each case by multiplying the coefficient by the cpm for the named gene and summing them. We then derived a single cutpoint for that new metric that best divided patients into high and low risk of decompensation using the surv_cutpoint function in R. Based on that, we then classified them into high and low risk groups to generate the Kaplan-Meier plot. We repeated this model building separately using the modules linked to for NAD (GO:0070403, GO:0070404 and GO:0019674), and ECM (GO:0030198) including the genes annotated to the respective GO or child terms using AnnotationDbi::select in R. We scaled and centered the model metrics to allow sensible comparison using Cox hazard ratios. As a benchmark, we used scaled and centered FIB-4 values (calculated from values +/- 180 days of the biopsy and not calculable on all cases) as an established prognostic metric.

AIFM2 inhibition experiments

For AIFM2 inhibition experiments we used FoP-Heps. Cells were exposed to OA/PA and PBMC co-culture as described above with or without 1 µM of the AIFM2 inhibitor iFSP1 (Hölzel; HY-136057) with or without 50 µM FAD. We assessed viability using the PrestoBlue assay according to manufacturer´s instructions and measured ROS and steatosis levels as described above.
